# Genomes of Two Flying Squid Species Provide Novel Sights into Adaptations of Cephalopods to Pelagic Life

**DOI:** 10.1101/2022.03.14.484290

**Authors:** Min Li, Baosheng Wu, Peng Zhang, Ye Li, Wenjie Xu, Kun Wang, Qiang Qiu, Jun Zhang, Jie Li, Chi Zhang, Jiangtao Fan, Chenguang Feng, Zuozhi Chen

**Author notes:** Corresponding author(s). (Chen Z), (Feng C). Equal contribution (Min Li, Baosheng Wu).

## Abstract

Pelagic cephalopods have evolved a series of fascinating traits, such as excellent visual acuity, high-speed agility, and photophores for adaptation to open pelagic oceans. However, the genetic mechanisms underpinning these traits are not well understood. Thus, in this study, we obtained high-quality genomes of two purpleback flying squid species (*Sthenoteuthis oualaniensis* and *Sthenoteuthis* sp.), with sizes of 5450 and 5651 Mb. Comparative genomic analyses revealed a common expansion of the S-crystallin subfamily *SL20-1* associated with visual acuity in the purpleback flying squid lineage and showed that evolution of high-speed agility for the species was accompanied by significant positive selection pressure on genes related to energy metabolism. These molecular signals might have contributed to the evolution of their adaptative predatory and anti-predatory traits. In addition, transcriptomic analysis provided clear indications of the evolution for the photophores of purpleback flying squids, *inter alia* that recruitment of new genes and energy metabolism genes may have played key functional roles in the process.

## Introduction

Cephalopods are a group of marine mollusk with remarkable morphology and behavior that play key ecological roles, are commercially important, and have been intensively studied [1]. Large populations of cephalopods inhabit depths ranging from shallow to abyssal [2, 3]. They are preyed upon by various apex predators (such as billfish, tuna, sharks, and cetaceans) that are generally the fastest and most efficient in the ocean [4-6]. In response to the predation, most cephalopods have developed keen visual acuity [7-9], strong muscles, and morphological traits [10] that enable them to evade danger rapidly. They also have high metabolic levels and hence constantly high levels of energy supplies for their muscles [11]. The continuous adaptations and counter-adaptations induced by interactions between prey and predators—the ‘arms race’, is one of the most intense forms of co-evolution. This ‘arms race’ is particularly pronounced between the pelagic cephalopods, *e.g*. cuttlefishes and squids, and their predators [12]. These cephalopods have extremely strong muscles with obliquely-striated, quickly-contractible, highly aerobic fibers, and morphological features, such as fins, that enable powerful swimming [10]. Some species of pelagic cephalopods have also repeatedly evolved photophores that assist escape [13], predation [14], and mating [15]. However, although it has long been accepted that the adaptative evolution has resulted in a series of fascinating traits in cephalopods [16-19], the genetic mechanisms involved are much less clear.

Common pelagic cephalopods include members of the Ommastrephidae (Teuthida, Decapodiformes) called flying squids, that can jump out of the water and in some cases glide more than 30 m in the air [20]. They can achieve the fastest recorded speed (∼ 8 m s^−1^) of any aquatic invertebrates [21, 22]. Important taxa with such capabilities include the purpleback flying squids (*Sthenoteuthis* spp.) [21, 23, 24]. These are the most abundant large squids in the tropical and subtropical Indo-Pacific ocean, found at depths from the surface to more than 600 m [2]. Moreover, the purpleback flying squids have a high degree of adaptation to their pelagic life and at least five morphological and ecological forms in terms of body size and possession or absence of photophores [2]. Thus, the purpleback flying squids are ideal models for studying the genetic mechanisms involved in the evolution of pelagic cephalopods’ adaptive traits.

In this study, we constructed high-quality genomes for two ‘forms’ of purpleback flying squids. One (*S. oualaniensis*) is the ‘medium’ or ‘typical form’, which has a dorsal mantel length (at maturity) of >120 mm and spherical photophores forming an oval patch in the anterodorsal mantle musculature [2, 25] (Figure S1). The other (*Sthenoteuthis* sp.) is the ‘dwarf form’, which is smaller and lacks the dorsal photophore patch (Figure S1). The dwarf form was previously treated as *S. oualaniensis*, but is now considered an undescribed species [26, 27]. Through comparative genomics analyses, we deeply profiled genomic features of these two purpleback flying squids and investigated molecular signals associated with adaptations of pelagic life in cephalopods, such as their excellent visual system, high behavioral flexibility, and photophore. The results suggest that genomes of these two purpleback flying squids are important resources that can facilitate research not only on cephalopods but also on adaptive evolution and molecular genetics more generally.

## Results and Discussion

### Assemblies and genomic characteristics

Using a combination of PacBio long reads, 10 × Genomics short reads, RNA sequencing, and Hi-C approaches, we obtained high-quality genomes of the two species of purpleback flying squids of the Ommastrephidae (*S. oualaniensis* and *Sthenoteuthis* sp.) (Tables S1–S4 and Figures S2–S3). The sizes of the genome assemblies were 5,450 and 5,651 Mb for *S. oualaniensis* and *Sthenoteuthis* sp., respectively (**Table 1**), close to the genome sizes estimated by K-mer analysis (Figure S4). The *S. oualaniensis* genome was assembled to chromosome level, with 95.72% of contigs anchored to the 46 chromosomes (**Figure 1**A). The contig and scaffold N50 values of the *S. oualaniensis* assembly are 1.52 Mb and 118.8 Mb, respectively, while the contig N50 value for the *Sthenoteuthis* sp. assembly is 1.3 Mb. The *S. oualaniensis* genome is the only squid genome that has been assembled to chromosome level (Table 1 and Figure 1A). GC contents of the acquired *S. oualaniensis* and *Sthenoteuthis* sp. genomes are 33.70 % and 33.00 %, respectively, similar to those of other relatives (Table S5). Results of BUSCO [28] assessment suggest that the *S. oualaniensis* and *Sthenoteuthis* sp. assemblies have 89.8% and 93.6% genomic integrity, respectively (Table 1). RNA-seq analysis of *S. oualaniensis* and *Sthenoteuthis* sp. yielded 389,954 and 410,364 transcripts with total lengths of 200–25,632 bp and 200–31,105 bp, respectively (Table S6).

**Table 1.** Summary of the assemblies and annotations of the two *Sthenoteuthis* species.

|  | *Sthenoteuthis oualaniensis* | *Sthenoteuthis* sp. |
| --- | --- | --- |
| Genome size (Mb) | 5,450 | 5,651 |
| Contig N50 (Mb) | 1.52 | 1.3 |
| Scaffold N50 (Mb) | 118.8 | - |
| GC content | 33.07% | 33.00% |
| Chromosome number | 46 | - |
| Gene number | 26,646 | 28,715 |
| BUSCOs for genomes | 89.4% | 93.6% |
| BUSCOs for genes | 90.4% | 94.3% |

**Figure 1.**
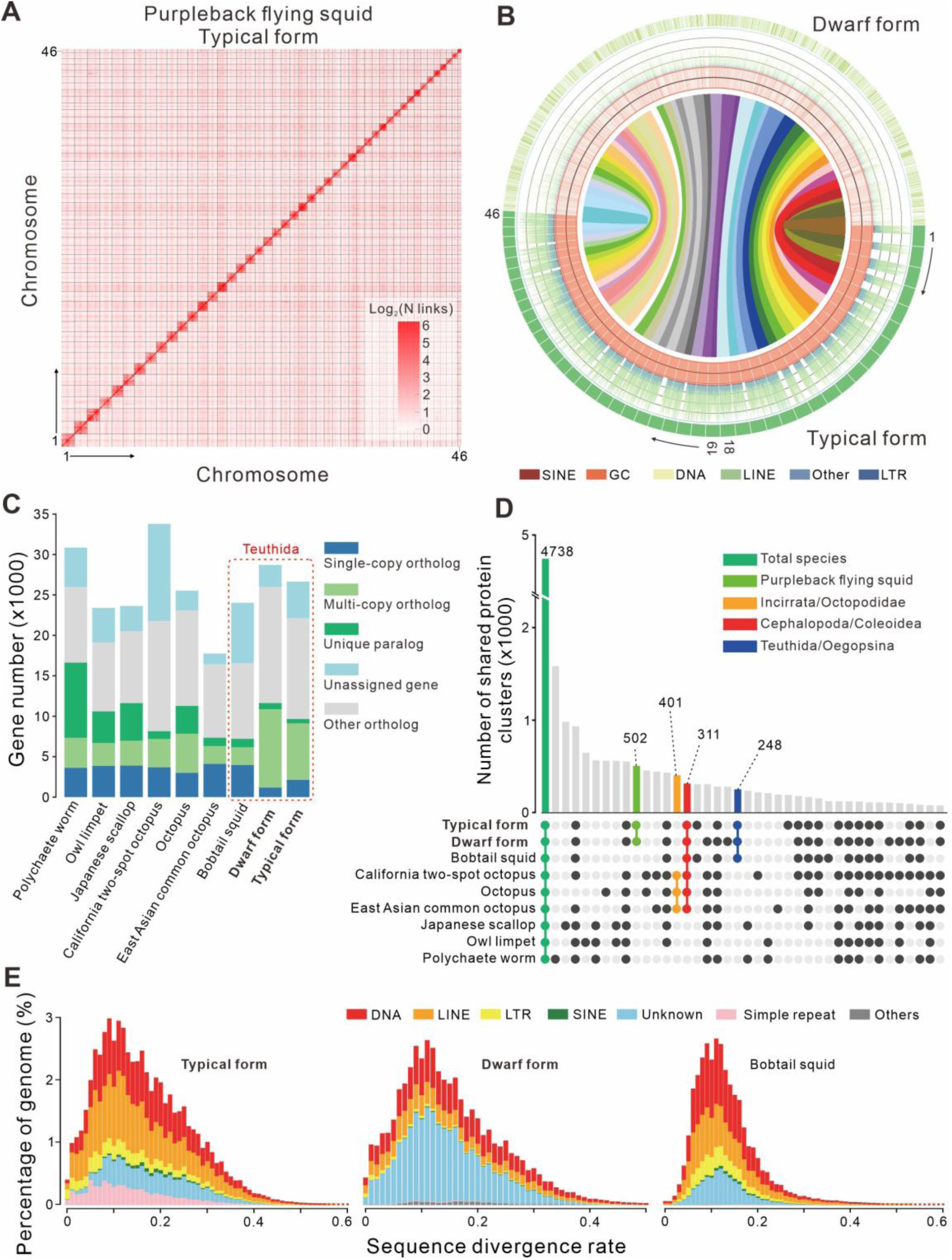
Genome assemblies and annotations of two *Sthenoteuthis* species. “Typical form” and “Dwarf form” of purpleback flying squids are referred to as *S. oualaniensis* and *Sthenoteuthis* sp., respectively. **A**. Genome-wide Hi-C map of the 46 pseudo chromosomes of *S. oualaniensis*. **B**. Syntenic comparison of *S. oualaniensis* and *Sthenoteuthis* sp. Numbers 1 to 46 refer to the chromosomes of *S. oualaniensis*. Each color block in the outermost layer represents a scaffold. Each line represents a syntenic block of five or more genes. Densities of specific kinds of TEs (ranging from 0 to 80%) were counted in 500 kb windows. **C**. Summary of gene clusters estimated from Orthofinder analysis, based on sequences of eight Mollusca species and one annelid worm. **D**. UpSet plot of gene families. **E**. Divergence distribution of TEs of species in Teuthida. Latin binomials of the listed species are as follows: Polychaete worm, *Capitella teleta*; Owl limpet, *Lottia gigantea*; Japanese scallop, *Mizuhopecten yessoensis*; California two-spot octopus, *Octopus bimaculoides*; Octopus, *Octopus minor*; East Asian common octopus, *Octopus vulgaris*; Bobtail squid, *Euprymna scolopes*.

Based on the high-quality assemblies, we predicted a total of 26,646 and 28,715 protein-coding genes in the genomes of *S. oualaniensis* and *Sthenoteuthis* sp., respectively (Table S5). Of these, 19,913 (74.73%) and 21,853 (76.10%) are supported by corresponding transcripts and 24,845 (93.24%) and 27,897 (97.13%) predictions could be functionally annotated using entries in at least one database (Figures S5–S6). Results of BUSCO analysis suggested that the assembly completeness of *S. oualaniensis* and *Sthenoteuthis* sp. were 89.4% and 93.6%, respectively, and the annotation completeness of them were 90.4 and 94.3%, respectively (Table 1 and Table S7). Basic metrics for the protein-coding genes of these two purpleback flying squids, including gene number/length, exon number/length, and codon usage, are close to those of other Mollusca (Table S5). Synteny analysis, based on the coding genes, identified 743 blocks with at least five syntenic genes between *S. oualaniensis* and *Sthenoteuthis* sp. and demonstrated the good colinearity between the two genomes, which also indicated that the two genomes were well assembled and annotated (Figure 1B). Finally, 22,162 (83%) *S. oualaniensis* genes and 25,991 (90%) *Sthenoteuthis* sp. genes were clustered into 16,235 gene clusters, with sizes close to those of other relatives (Figure 1C and 1D).

Repeats analysis indicated that close to half of the genomes of both *S. oualaniensis* and *Sthenoteuthis* sp. are composed of repetitive sequences: 53.90% and 42.37%, respectively (Tables S8-S9), which is comparable to that of other cephalopods [29, 30]. Identification and classification of the TEs of the two genomes (Figure 1E and Tables S8–S9) showed that DNA and LINE were the two most abundant types of TEs in the *S. oualaniensis* genome (accounting for 25.47 and 22.05% of the total, respectively), but only accounted for 14.13% and 10.00% of the TEs, respectively, in the *Sthenoteuthis* sp. genome. There was a markedly smaller proportion of simple tandem repeats in the *S. oualaniensis* genome (6.33%) than in the *Sthenoteuthis* sp. genome (25.86%) (Tables S8–S9). Thus, despite the very close relationship of the two purpleback flying squid species, they might have distinct patterns of TE activity.

### Phylogenetic status and species validity of two purpleback flying squids

ML gene tree and species tree analyses based on 334 one-to-one genes yielded consistent topologies (Figure S7). The Octopodiformes and Decapodiformes species each clustered into monophyletic lineages, which apparently diverged around 366.5 Ma (**Figure 2** and Figure S7). The two purpleback flying squid species are the most closely related of the included taxa, having diverged at approximately 41.0 Ma. These two forms of purpleback flying squids, *S. oualaniensis* and *Sthenoteuthis* sp., were considered to be members of the same species for a long time, but this was recently questioned [26, 27]. Morphological studies provided evidence that *Sthenoteuthis sp*. was distinguishable from *S. oualaniensis* concerning external features, including variables of the head, carcass, arms, as well as the shape and size of fins [31, 32]. Our results corroborate the conclusion that *Sthenoteuthis* sp. should be regarded as a distinct species, as we detected clear differences in their genomes and derived a substantial divergence time.

**Figure 2.**
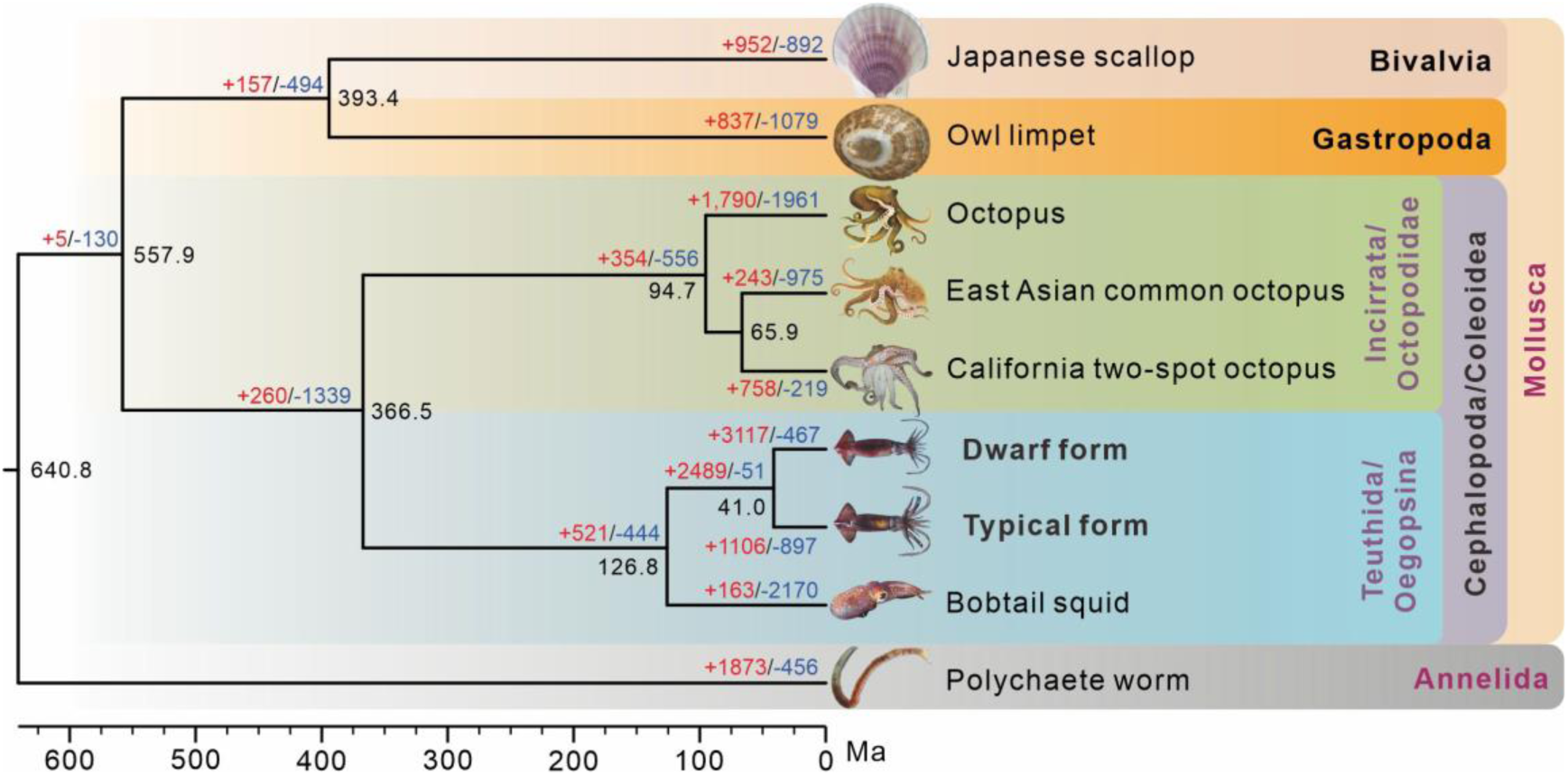
Coalescence tree of eight Mollusca species and one annelid worm based on 334 single-copy orthologues. Estimated divergence times and expanded/contracted gene families are marked at the nodes. Red, blue and black numbers indicate expanded gene families, contracted gene families, and estimated divergence times (million years ago, Ma), respectively.

### Excellent vision of purpleback flying squids

Keen vision is regarded as a major product of evolutionary arms races in squids [7], which strongly contributes to the ability of purpleback flying squids (and others) to avoid predation and catch prey even in dim conditions [1, 7, 9]. Important components of their eyes include several soluble crystallins that play key roles in maintaining the transparency and optical clarity of the lens [33]. In particular, S-crystallins are present in lenses of many cephalopods, and have refractive properties that strongly contribute to good vision (and hence cephalopod survival) in poor light [34, 35]. S-crystallins are even claimed to provide a “perfect medium”, forming gels of varying density, in the spherical lenses of cephalopods [35-37].

Cluster analysis detected 2489 gene families that appear to have expanded in both purpleback flying squid species (under the criterion of both Family-wide and Viterbi *P*-values < 0.01; Figure 2 and Figures S8–S9). The subfamily *SL20-1* of S-crystallins is the most significantly expanded, with 39 and 99 gene members in *S. oualaniensis* and *Sthenoteuthis* sp., respectively (**Figure 3**A and 3B). Transcriptomic analysis revealed that most of these expanded *SL20-1* genes were only highly expressed in eyes (Figure 3C and Figure S10), clearly indicating that expansion of this subfamily played an important role in the emergence of purpleback flying squids’ excellent vision.

**Figure 3.**
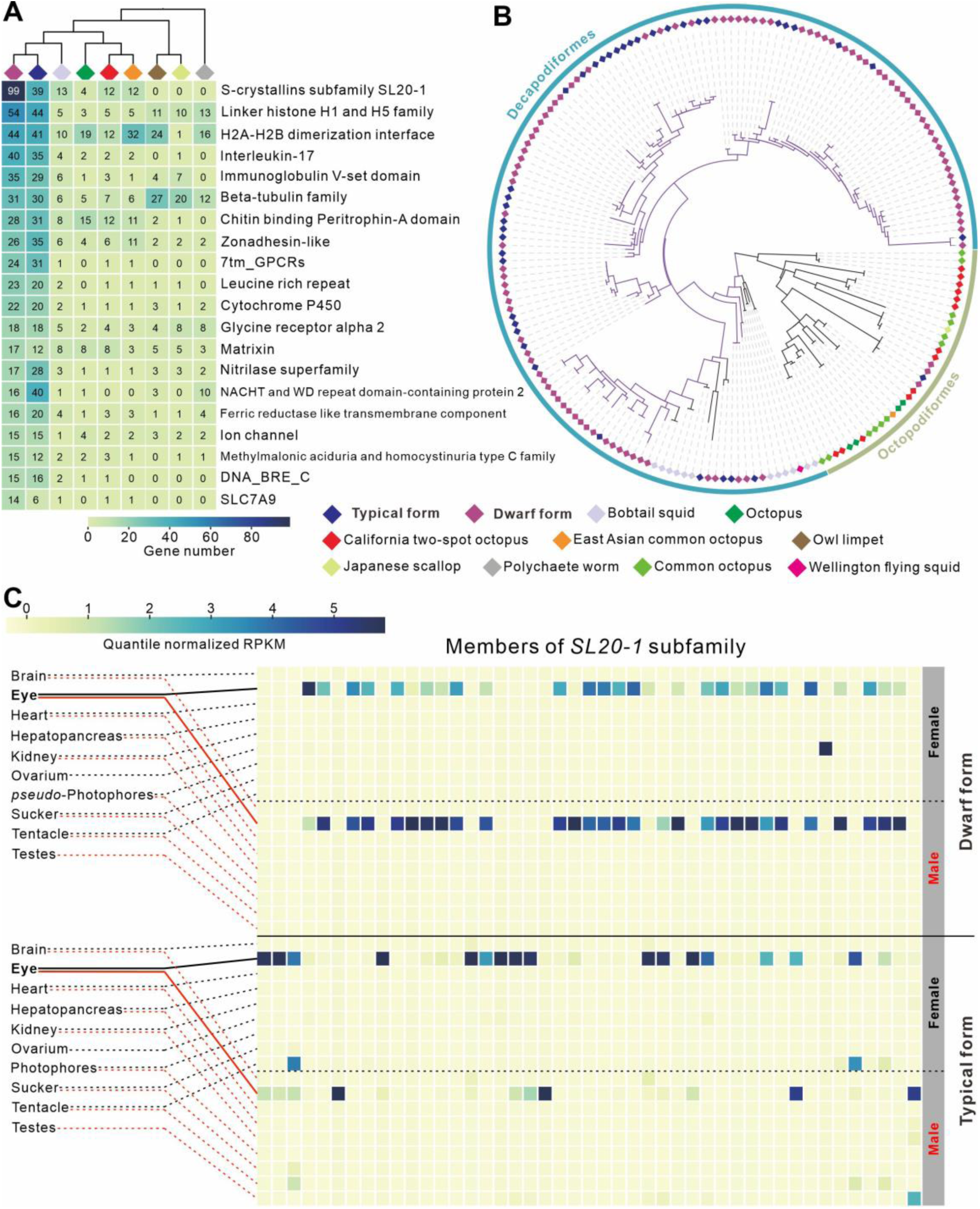
Expanded gene families in the *Sthenoteuthis* lineage. **A**. The top 20 expanded gene families. **B**. Unrooted maximum-likelihood tree of the massive expansion of the S-crystallin subfamily *SL20-1*, which has 39 and 99 gene members in *S. oualaniensis* and *Sthenoteuthis* sp., respectively. **C**. Expression patterns of coexisting gene members of the *SL20-1* subfamily in *S. oualaniensis* and *Sthenoteuthis* sp. Only those expressed gene copies were shown. These gene IDs were detailed in Figure S10. Most of the significantly expanded genes were highly expressed in the eyes. Latin binomials of the listed species are shown in the legend of Figure 1 except Comment octopus, *Octopus sinensis*; Wellington flying squid, *Sepia pharaonis*.

### High behavioral flexibility

The mobility of organisms strongly affects their predatory and anti-predatory abilities, and thus their evolutionary fitness [38]. Purpleback flying squids are highly successful in these terms, as they are the fastest and most mobile aquatic invertebrates, with very high metabolic levels that enable powerful output at all times [11, 24].

Our analysis identified 66 PSGs in the *Sthenoteuthis* lineage (with FDR-adjusted *P*-values < 0.01, Table S10). GO enrichment analyses (FDR <0.01) indicated that 23, 42, and 30 of these PSGs are associated with cellular components, molecular functions, and biological processes, respectively (Figures S11–S12). The PSGs involved in energy metabolism have apparently been under significant evolutionary selection pressure (FDR-adjusted *P*-values < 0.01). “Phosphoglycerate kinase activity” (GO:0004618, *e.g. pgk1*) and “[2Fe-2S] cluster assembly” (GO:0044571, *e.g. IscS*) were the two most significant GO terms (FDR < 0.05; Tables S11–S12). The “quinone binding” category (GO:0048038, e.g. *ndufa6* and *ndufa7*) was also significantly enriched (FDR < 0.05; Table S11). Transcriptomic analysis demonstrated that these genes are highly expressed in all the investigated tissues, implying that they play important roles (**Figure 4**A).

**Figure 4.**
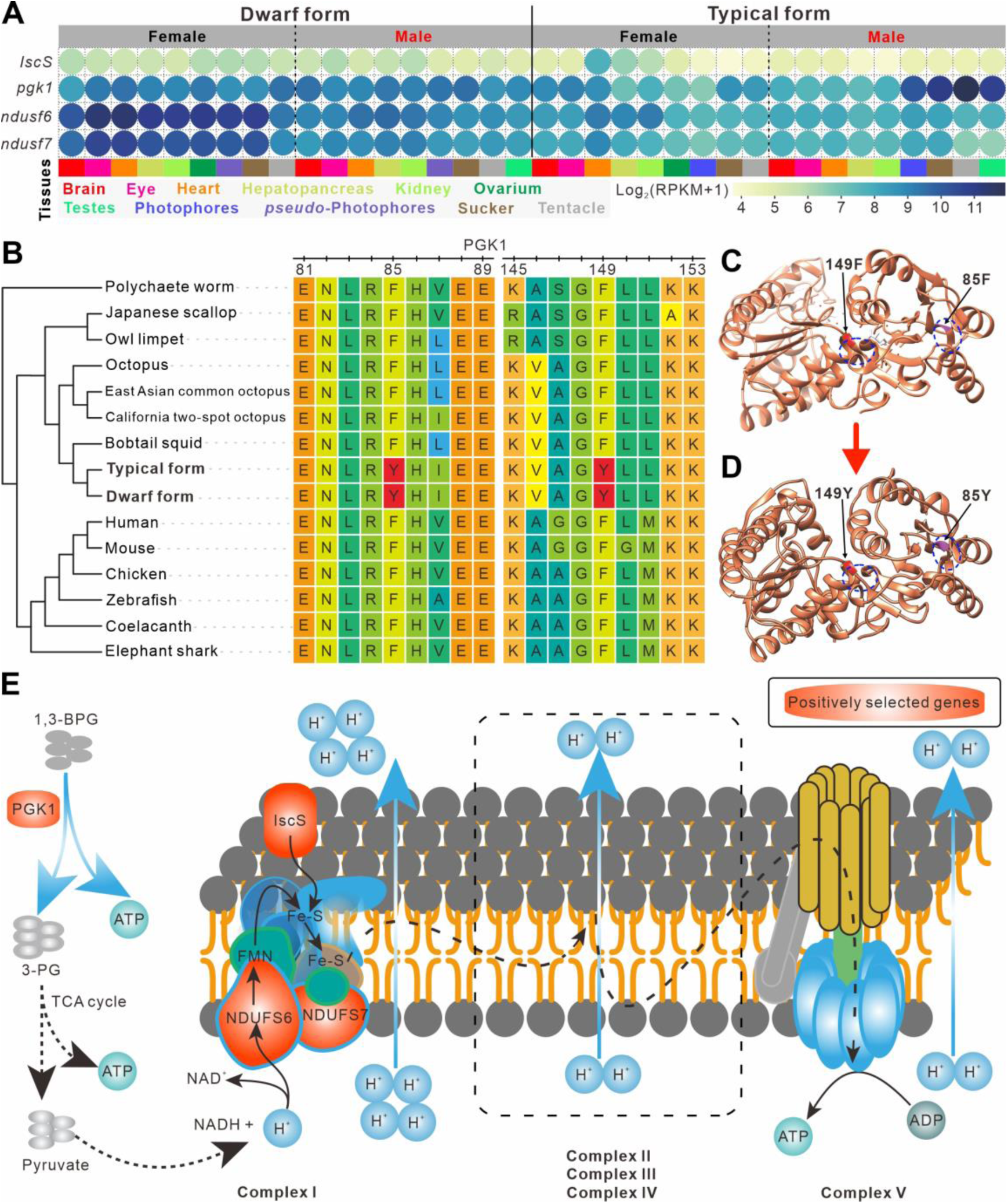
Diagram of positively selected genes associated with energy metabolism. **A**. Expression patterns of four positively selected genes involved in energy metabolism that are highly expressed in all investigated tissues. **B**. Positively selected signals in two extremely conserved regions of *pgk1* gene. **C**. Three-dimensional structure of mouse PGK1 protein downloaded from the PDB database. Substructures of 85F and 149F are highlighted. **D**. Three-dimensional structure simulated by a homologous approach of mouse PGK1 with F85Y and F149Y substitutions. Structures adjacent to the substitute sites in panels **C** and **D** are signaled by the blue dashed circles. **E**. Schematic diagram of the glycolysis pathway and respiratory electron chain.

Phosphoglycerate kinase 1 (PGK1), the first ATP-generating enzyme in the glycolytic pathway, both directly generate ATP and indirectly supply fuels for the mitochondrial electron respiratory chain [39, 40]. PAML analysis detected positive selection signals at two extremely conserved loci of the *pgk1* gene (F85Y and F149Y, Figure 4B). Simulations showed that these two F-to-Y substitutions affect the structure of PGK1 protein resulting in the conversion of the original helix to a loop (Figures 4C and 4D). Thus, these substitutions were presumably highly important for purpleback flying squids. Similarly, a positive selection signal at a conserved locus was detected in the *IscS* gene (Y67H, Figure S13). *IscS* participates in the synthesis of multiple iron-sulfur (Fe-S) proteins and the formation of Fe-S clusters in Complex I (NADH: ubiquinone oxidoreductase; Figure 4E) [41]. Knockdown of the *IscS* gene leads to a decrease in mitochondrial activity [41, 42]. We also found positive selection signals for genes encoding two important subunits of Complex I, *ndufa6* and *ndufa7* (Table S10), which are essential for the catalytic activity of the complex [43, 44]. Complex I plays a key role in ATP synthesis driven by the mitochondrial electron respiratory chain [45](Figure 4E). Therefore, the positively selected genes mentioned above are likely to promote ATP synthesis, which is important for the maintenance of high metabolic levels and the high behavioral flexibility of purpleback flying squids. Furthermore, the presence of these positively selected sites in the pelagic *Architeuthis dux* (Figure S14), which also possess high metabolic levels [2, 30], implies the importance of these selection signals for pelagic cephalopods. But we should note that these results come from a small gene pool and therefore have some limitations.

### Photophore transcriptome

Bioluminescence is a common feature of cephalopods, especially pelagic species [14, 46]. At least 63 genera of squid and cuttlefish have repeatedly evolved photophores that play important roles in defense, predation, and communication [13-15]. One of the species included in this study, *S. oualaniensis*, has a dorsal photophore patch, but not *Sthenoteuthis* sp. (Figure S1). Therefore, they are ideal models for studying the evolution of photophores. PCA showed that *S. oualaniensis* photophores clustered with the *pseudo*-photophores (‘muscle tissue’ corresponding to the position of the photophore of *S. oualaniensis*) of *Sthenoteuthis* sp. (**Figure 5**A) and highly expressed genes of photophores had similar expression patterns (Figure 5B). Therefore, the *pseudo*-photophores of *Sthenoteuthis* sp. and photophores of *S. oualaniensis* seem to be homologous organs, and the *pseudo*-photophores may have some essential functions similar to those of the photophores.

**Figure 5.**
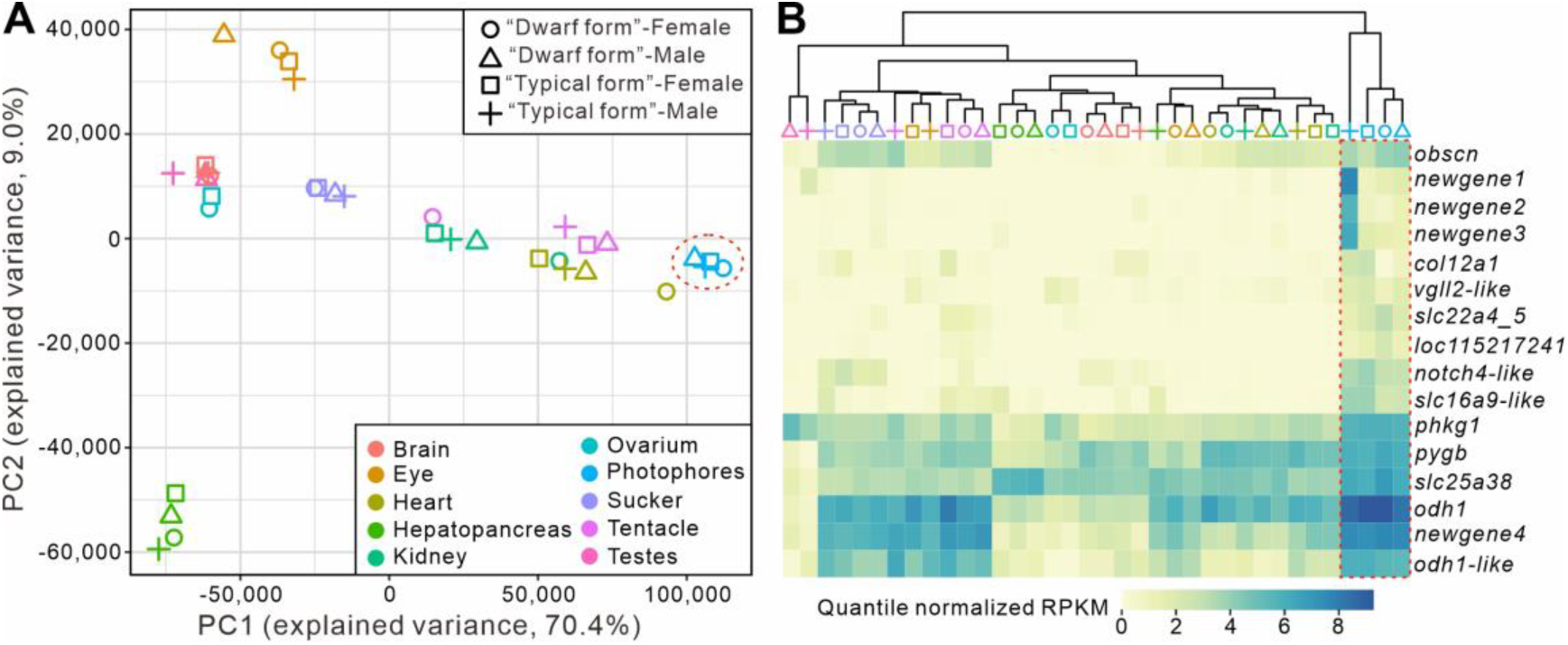
Results of photophore transcriptome analysis. **A**. PCA plot of 36 RNA-seq samples based on expression profiling of 23,082 orthologs. *Pseudo*-photophores of *Sthenoteuthis* sp. and photophores of *S. oualaniensis* clustered together (red dotted circle). **B**. Whole-tissue expression patterns of 16 highly expressed genes in the four “photophore” tissues.

Previous studies based on gene expression profiles have suggested that the massive recruitment of pre-existing gene modules plays an important role in the formation of cephalopod photophores [47, 48]. Here we specifically investigate highly expressed genes in the “photophores” of these two squids, which represent genetic factors associated with the essential function of this organ. Among the 16 highly expressed genes, four were new genes (*newgene1*-*4*; Figure 5B). Although functional assignations of these genes were not supported by homologs in public databases, their specific expression patterns suggest that they might participate in the formation and some of the basic functions of luminophores. In addition, four of the highly expressed genes (*phkg1, pygb, odh1*, and *obh1*-like) are energy-metabolism-related genes. All of these genes are involved in glucose metabolism. Both *phkg1* and *pygb* are glycogen phosphorylases that regulate the catabolism of glycogen and provide glucose used in glycolysis [49, 50]. Octopine dehydrogenase (coded by *odh1*), mainly found in mollusks [51], has a major function similar to that of lactate dehydrogenase, providing an important reducing agent for the glycolytic process [52]. These observations imply that these new genes and recruited energy-metabolism-related genes might provide important support for the photophores of purpleback flying squids.

## Conclusion

In this study, we generated high-quality genomes of the two purpleback flying squid species. Comparative genomics analyses indicated that expansion of the S-crystallin subfamily *SL20-1* and locus variation and expression patterns of genes related to energy metabolism are associated with adaptations of purpleback flying squids (such as excellent vision, high behavioral flexibility, and photophore) that have played important roles in the ‘arms race’ and other pelagic adaptations among marine organisms. Moreover, the study supports the validity of treating *Sthenoteuthis* sp. as a separate species at the genomic level. These findings advance our understanding of the genetic basis of pelagic cephalopods associated with predatory and anti-predatory traits and suggest that the two genomes could be important resources in studies not only of cephalopods, but also co-evolution, bioluminescence, and other broader aspects of molecular genetics.

## Materials and methods

### Sampling and sequencing

Squid samples were caught by a commercial fishing vessel using a lit falling net at night in the South China Sea. One male and one female individual of both medium (*S. oualaniensis*) and dwarf (*Sthenoteuthis* sp.) purpleback flying squids were collected for sequencing. Muscles from the males were used for DNA extraction and genomic library preparation. A PacBio Sequel device was used for sequencing the long reads. Short-insert paired-ends libraries were prepared and sequenced according to the Illumina sequencing protocol. Sample indexing and partition barcoded libraries were prepared using a Chromium Genome Reagent kit (Cat. PN-120229, 10× Genomics, USA) and sequenced by an Illumina HiSeq X-Ten system for Hi-C analysis of *S. oualaniensis*. To explore gene expression patterns of the species and aid gene annotation, RNA was extracted from nine tissues—tentacle, brain, eye, heart, kidney, sucker, hepatopancreas, ovarium/testes, and photophore (for *Sthenoteuthis* sp., muscle tissue corresponding to the position of the photophore of *S. oualaniensis* were obtained)—from each of the four *S. oualaniensis* and *Sthenoteuthis* sp. individuals for library preparation and sequenced using the Illumina HiSeq 2000 platform.

### Estimation of genome sizes of the two *Sthenoteuthis* species

The genome sizes of the two *Sthenoteuthis* species were estimated by *k*-mer analysis using filtered Illumina reads. We used SOAPec v2 [53] to estimate the distribution of 17-mer depth, then estimated the genome size from the total base and peak values of 17-mer depth.

### Genome assembly

Based on the estimated genome sizes, we first assembled genomes of the two purpleback flying squid species to contig level with wtdbg2 v2.4.1 [54] and standard parameters. The Arrow algorithm was used to polish the two draft genomes with the filtered PacBio reads. Then the filtered short paired-end reads were aligned to the draft genome by BWA-MEM v.0.7.12-r1039 [55] with standard parameters, and Pilon [56] was used to further polish the genomes in two rounds using the sorted bam files. Finally, the genome assembly of *S. oualaniensis* was anchored with the Hi-C reads by 3D-DNA [57] and Juicer v1.5 [58]. To improve the quality of the chromosome assembly, we used Juicebox Assembly Tools [59] to remove potential assembly errors. BUSCO v3.02 [28] with the “metazoa_odb9” library was used to evaluate the completeness of the two assemblies.

### Repetitive sequence annotation

After obtaining a high-quality genome assembly we used a combination of *de novo* and homologous predictions to annotate repetitive sequences, including tandem repeats and transposable elements (TEs). Firstly, tandem repeat finder v4.07 [60] was used to scan the tandem repeat elements with the parameter settings “2 7 7 80 10 50 500 -d -h -ngs”. Then we used RepeatModeler v1.0.8 [61] to build a *de novo* repeat library, and RepeatMasker v3.3.0 [62] to detect homologous repeat elements. After integrating the results of *de novo* and homologous predictions, Jukes-Cantor distances were calculated and the R8s algorithm [63] was used to calculate rates of evolution from them.

### Protein-coding gene prediction

A combination of *ab initio*, homologous, and transcript-based gene predictions was used to integrate the two genomes. The gene prediction pipeline was as follows. First, AUGUSTES v3.2.1 [64], GlimmerHMM v3.02 [65], and GeneID v1.4 [66] were used for *de novo* gene prediction. Second, we downloaded the non-redundant proteomes of *Lottia gigantea* (GCF_000327385.1), *Mizuhopecten yessoensis* (GCF_002113885.1), *Octopus bimaculoides* (GCF_001194135.1), *Euprymna scolopes* (GCA_004765925.1), *O. minor* (http://dx.doi.org/10.5524/100503), and *O. vulgaris* (GCA_003957725.1) for homologous gene prediction. We used TBLASTN v2.9 [67] to align the proteomes of the six relatives to the two purpleback flying squid genomes and extended 10,000 bp in both directions from the start and end of every TBLASTN hit. Then, all non-redundant transcripts of all tissues were aligned to the genome with TBLASTN and a 1000 bp extension was applied. Genewise v2.4 [68] was then used to resolve the gene structure according to the above hits. Next, we integrated results of the *ab initio*, homologous, and transcript-based predictions with 1:4:5 weights using EvidenceModeler [69]. Finally, for further functional annotation of these two gene sets, we scanned public databases, including Swiss-Prot, KOG, Nr, KEGG, Gene Ontology (GO), and Pfam to detect the best matches using Interproscan v5 [70].

### Gene family clustering analysis

In addition to the two predicted gene sets and six Mollusca species above, we downloaded the genome of *Capitella teleta* (GCA_000328365.1) as an outgroup. Proteomes of these nine species were subjected to an all-vs-all BLAST search (e-value ≤ 1e^-6^) then clustered by OrthoFinder [71] with default parameters. The shared gene clusters were visualized by the R package UpSetR [72]. The expanded and contracted gene families were investigated by CAFÉ v4.0.1 [73] using the result of the clustering analysis under the criterion of both Family-wide and Viterbi *P*-values < 0.01.

### Phylogeny and divergence time estimation

Based on the cluster analysis of the above nine species, the protein-coding sequences and corresponding codon sequences of 334 one-to-one homologous genes were picked out and aligned using MAFFT v7 [74]. The bad alignments were removed by trimAl [75]. Finally, we used RAxML v8.2.4 [76] with “-m GTRGAMMA -f a -x 271828 -N 100 -p 54321” parameter settings to construct phylogeny trees and ASTRAL [77] to infer a species tree. MCMCtree in the PAML package [78] was used to estimate divergence times in conjunction with two softbound calibration points from www.timetree.org: *O. bimaculoides* - *C. teleta* (585 - 679 Ma) and *O. bimaculoides* - *L. gigantea* (531 - 582 Ma).

### Synteny between the two *Sthenoteuthis* species

To evaluate the conservation and quality of the two assemblies of *Sthenoteuthis* species, we used the *S. oualaniensis* assembly as a reference and *Sthenoteuthis* sp. assembly as the query in alignment analysis by LAST v942 [79] with the -E0.05 parameter setting. We also calculated densities of repetitive elements (DNAs, LINEs, TRFs, SINEs, and Others) and GC content in 500,000 bp windows of the genomes. Finally, the results of these analyses were integrated into a circular layout by CIRCOS v0.69 [80].

### Positive selection analysis

To evaluate the evolutionary pressure on *Sthenoteuthis*, we used the one-to-one homologous genes of the nine species listed above (see section 2.6 *Gene family clustering analysis* for details) to identify positively selected genes (PSGs). We aligned codons of all the one-to-one homologous genes by PRANK v140603 [81] with “-codon -f=fasta”. All the gaps generated by alignments were removed by Gblocks v0.91b [82] with “-t=c”. Then we used an in-house Perl script to convert the aligned sequence to PAML format for use in PAML analysis. Finally, PAML 4.9i [78] was used to analyze the selection pressure on each gene with the maximum-likelihood method under the branch-site model. A species tree constructed from ASTRAL analysis was used as the input tree. The two *Sthenoteuthis* species were selected as foreground and the other seven species as background. The significance of the alternative model (estimated omega) against the null model (fixed omega) was assessed by likelihood ratio tests (LRTs), in which twice the log-likelihood difference (2DL) values were calculated and compared to a chi-squared distribution. Genes with a *P*-value ≤ 0.01 (with false discovery rate, FDR, correction), with at least one site under positive selection with a Bayes Empirical Bayes (BEB) posterior probability > 0.8, were identified as candidate PSGs.

### Protein structure simulation

A homology-based approach was used to simulate structures of proteins encoded by PSGs. We first sought matches to the *S. oualaniensis* PGK1 protein sequence in the PDB database (https://www.rcsb.org/) and selected the hit with the highest score as a potential template for simulation of the PGK1 protein structure. The corresponding positively selected sites (F85 and F149) of mouse PGK1 protein were replaced by those of *Sthenoteuthis* (85Y and 149Y). Then, the modified sequence was submitted to Phyre2 [83] for structure simulation. Finally, the Phyre2 result with the highest score was selected as the final structure and visualized by UCSF Chimera [84].

### Transcriptomic analysis

Raw reads obtained from RNA sequencing of the nine mentioned tissues of the four sampled individuals were filtered using fastp [85] with default parameters. The low-quality reads were removed by Sickle v1.33 (https://github.com/najoshi/sickle) with default parameters except for the “pe” setting. The cleaned reads were mapped to the reference genome with Hisat2 [86]. Then Trinity [87] was used to assemble these reads into transcripts. Next, we used TransDecoder [88] with default parameters to predict gene structures of the transcripts and CD-HIT [89] to remove redundant predictions. The numbers of reads and RPKM (Reads Per Kilobase of transcript per Million mapped reads) values for all genes in the 36 tissues were calculated by StringTie v2.1.4 [90] using the output of Hisat2 [86] analysis. Each gene with an RPKM value greater than 1 was considered as a validly expressed gene. The *Tau* value of genes in all tissues was calculated using an in-house Perl script. Genes expressed in the photophore with *Tau* values ≥ 0.8 and higher RPKM values than in other tissues were regarded as being highly specifically expressed in the photophore. All the RNA-seq data for the 36 tissues were subjected to Principal Component Analysis (PCA), using normalized and logarithmically transformed RPKM values. Seaborn [91] was used to cluster and visualize clusters of the highly expressed genes in the photophore. Those genes shared only by the two *Sthenoteuthis* species and supported by transcripts were identified as new genes. To investigate the expression pattern of expanded genes of the SL20-1 subfamily across organs, RNA sequencing reads of both species were mapped to the genomes of *Sthenoteuthis* sp. and subsequently assessed by RPKM values.

## Data availability

The genome and annotation of these two purpleback flying squid species and raw sequence data and acquired RNA-seq data were deposited in the China National GeneBank (CNGB, https://db.cngb.org/cnsa) under accession number CNP0001958 (https://db.cngb.org/cnsa/project/CNP0001958/reviewlink/).

The raw sequencing data (accession ID CRA004867, at https://ngdc.cncb.ac.cn/gsa/s/D7aN3eh2) and the genomes and annotations (accession ID GWHBECU00000000 and GWHBFHL00000000 for *S. oualaniensis* and *Sthenoteuthis* sp., respectively, at https://bigd.big.ac.cn/gwh) reported in this study were deposited in the Genome Sequence Archive and Genome Warehouse, respectively, in National Genomics Data Center, Beijing Institute of Genomics (China National Center for Bioinformation), Chinese Academy of Sciences.

## CRediT author statement

**Min Li:** Conceptualization, Validation, Formal analysis, Investigation, Project administration, Writing – original draft. **Baosheng Wu:** Formal analysis, Data Curation, Visualization, Writing – original draft. **Peng Zhang:** Validation, Resources, Investigation, Writing - Review & Editing. **Ye Li:** Formal analysis, Data Curation, Visualization. **Wenjie Xu:** Formal analysis, Data Curation, Visualization. **Kun Wang:** Writing - Review & Editing. **Qiang Qiu:** Writing - Review & Editing. **Jun Zhang:** Resources, Writing - Review & Editing. **Jie Li:** Resources, Investigation. **Chi Zhang:** Methodology, Software, Data Curation. **Jiangtao Fan:** Resources, Writing - Review & Editing. **Chenguang Feng:** Conceptualization, Methodology, Validation, Project administration, Writing – original draft. **Zuozhi Chen:** Supervision, Project administration, Funding acquisition, Writing - Review & Editing. All authors read and approved the final manuscript.

## Acknowledgments

This study was supported by a Guangdong Major Project of Basic and Applied Basic Research grant (No. 2019B030302004), the National Key Research and Development Program of China (grant no. 2018YFC1406502), the Key Special Project for Introduced Talents Team of Southern Marine Science and Engineering Guangdong Laboratory (Guangzhou) (grant no. GML2019ZD0605), the Financial Fund of the Ministry of Agriculture and Rural Affairs of China (grant no. NFZX2018), the Central Public-interest Scientific Basal Research Fund, CAFS of China (grant nos. 2019HY-XKQ03, 2020TD05, and 2021SD18), China Postdoctoral Science Foundation (2021M693342); Hubei Postdoctoral Innovation Post Project, the 1000 Talent Project of Shaanxi Province, and Research Funds for Interdisciplinary subject of Northwestern Polytechnical University, China (grant no. 19SH030408). We gratefully acknowledge colleagues at BGI-Shenzhen for data analyses and Dr. Shuai Zhang from SCSFRI, CAFS for collecting references and reviewing the manuscript.

## Supplementary material

**Figure S1 Morphology and dissection of the two *Sthenoteuthis* species** “Typical form” and “Dwarf form” of purpleback flying squids are referred to as *S. oualaniensis* and *Sthenoteuthis* sp., respectively. **A**. Morphology and the dorsal photophore patch of the “Typical form”. **B**. Morphology and the corresponding position of the photophore (pseudo-photophore) of the “Dwarf form”. **C**. Dissection of the female individual. **D**. Dissection of the male individual.

**Figure S2 Distribution of GC contents versus the sequencing depth for *S. oualaniensis***

The horizontal and vertical coordinates indicate the GC contents and sequencing depth respectively with 10 kb as a window. The GC contents of most windows were around 20%–50%, indicating the high-quality raw reads without significant contamination.

**Figure S3 Sequencing depth distribution for *S. oualaniensis***

The horizontal coordinate indicates the reads depth of PacBio reads. The vertical coordinate indicates the cumulative ratio of base depth. The blue line indicates the reads depth, the yellow line indicates the accumulation of all the reads depth.

**Figure S4 GenomeScope profiles for *S. oualaniensis* were estimated**

The result from jellyfish v2.2.6 with a *k*-mer size of 17 was used as the input file of GenomeScope v1.0. The second peak should be the major peak and has an estimated genome size of about 5.8 Gb. This is suggestive of the high heterozygosity of the species.

**Figure S5 Venn diagram of *S. oualaniensis* to show the well-aligned gene sets from a different database, including NR, InterPro, KEGG, SwissProt, and Trembl database**

**Figure S6 Venn diagram of *Sthenoteuthis* sp. to show the well-aligned gene sets from a different database, including NR, InterPro, KEGG, SwissProt, and Trembl database**

**Figure S7 Estimation of phylogeny (A.) and divergence times (B.) among purpleback flying squid species and their relatives**

The divergence time was indicated by the blue number near each node, while the 95% CI was indicated by the blue brackets and its specific value was colored in blue under the corresponding divergence time. Two softbound calibration time points had been applied: *Octopus bimaculoides* - *Capitella teleta* (585 - 679 MYA) and *Octopus bimaculoides* - *Lottia gigantea* (531 - 582 MYA).

**Figure S8 Top 20 KEGG enriched pathway of the significantly expanded gene families for both *Sthenoteuthis* species**

**Figure S9 The significantly enriched GO terms of the expanded gene families for both *Sthenoteuthis* species**

**Figure S10 Expression patterns of coexisting gene members of the *SL20-1* subfamily in *S. oualaniensis* and *Sthenoteuthis* sp**.

Only those expressed gene copies were shown. Gene IDs and tissue information correspond to Figure 3C. Most of the significantly expanded genes were highly expressed in the eyes.

**Figure S11 Classification of the three major GO function annotations for the 66 PSGs of the two *Sthenoteuthis* lineage**

**Figure S12 The significantly enriched KEGG pathway of the positively selected genes for both *Sthenoteuthis* species**

**Figure S13 Sequence alignments of *IscS* for the two *Sthenoteuthis* and other seven species from Mollusca and Annelida**

**Figure S14 Positive selection sites for genes *PGK1* (Figure 4B) and *IscS* (Figure S13) shared by the two *Sthenoteuthis* were also detected from the giant squid (*Architeuthis dux*)**.

**Table S1 PacBio reads statistics of the two *Sthenoteuthis* species**

**Table S2 Statistics of NGS sequencing**

**Table S3 Statistics of 10x sequencing**

**Table S4 Statistics of Hi-C sequencing**

**Table S5 Statistics of the predicted gene sets for two *Sthenoteuthis* species and the comparison with other relatives**

**Table S6 The sequencing reads information used for transcriptomes assembly from nine tissues**

**Table S7 The predicted genome completeness of the two *Sthenoteuthis* species with metazoa_odb10**

**Table S8 Transposable elements information for the genome of *S. oualaniensis***

**Table S9 Transposable elements information for the genome of *Sthenoteuthis* sp**.

**Table S10 66 positively selected genes for the purpleback flying squids’ clades with *P* <0.01**

**Table S11 Terms from the Function Ontology with FDR < 0.05 for the 66 PSGs of the two *Sthenoteuthis* lineage. The top 20 terms were listed**

**Table S12 Terms from the Process Ontology with FDR < 0.05 for the 66 PSGs of the two *Sthenoteuthis* lineage. The top 20 terms were listed**.

