## Supplementary materials for "Genomes of Two Flying Squid Species Provide Novel Sights into Adaptations of Cephalopods to Pelagic Life"

### Abstract

Pelagic cephalopods have evolved a series of fascinating traits, such as excellent visual acuity, high-speed agility, and photophores for adaptation to open pelagic oceans. However, the genetic mechanisms underpinning these traits are not well understood. Thus, in this study, we obtained high-quality genomes of two purpleback flying squid species (*Sthenoteuthis oualaniensis* and *Sthenoteuthis* sp.), with sizes of 5450 and 5651 Mb. Comparative genomic analyses revealed a common expansion of the S-crystallin subfamily *SL20-1* associated with visual acuity in the purpleback flying squid lineage and showed that evolution of high-speed agility for the species was accompanied by significant positive selection pressure on genes related to energy metabolism. These molecular signals might have contributed to the evolution of their adaptative predatory and anti-predatory traits. In addition, transcriptomic analysis provided clear indications of the evolution for the photophores of purpleback flying squids, *inter alia* that recruitment of new genes and energy metabolism genes may have played key functional roles in the process.

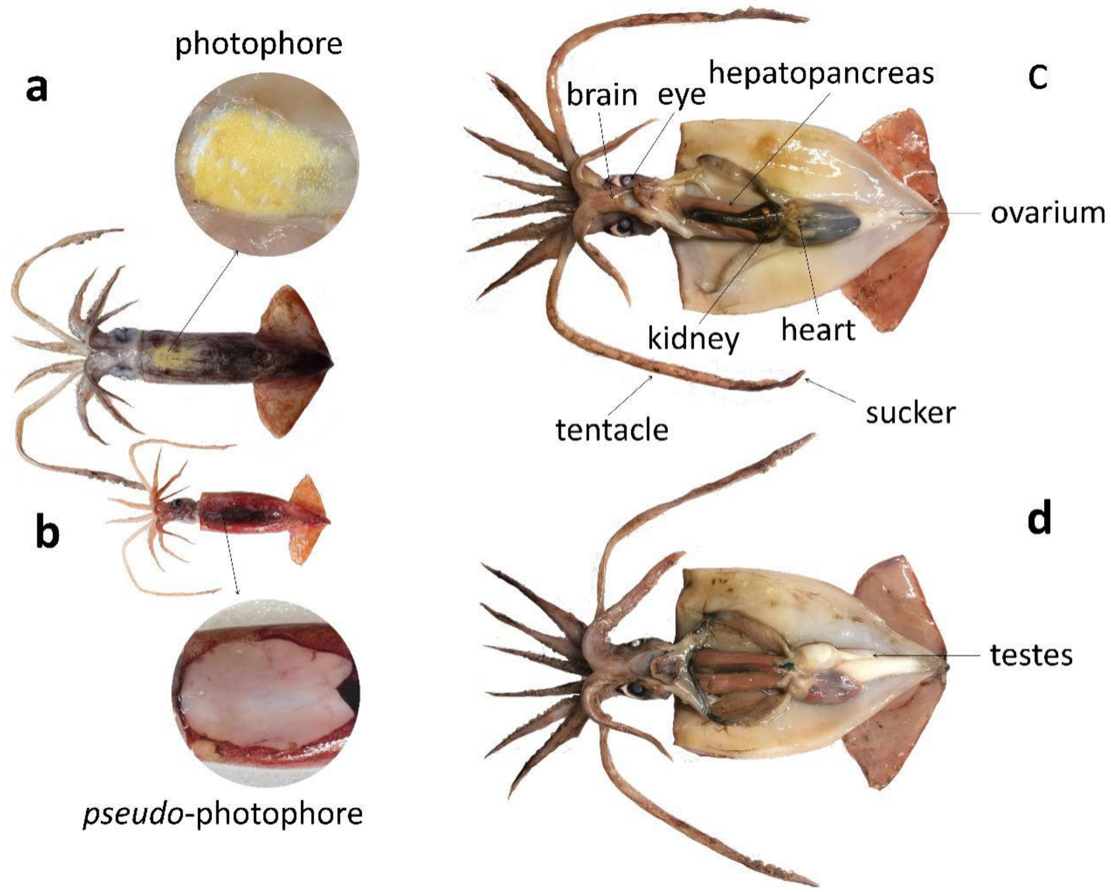


**Supplementary Figure S1 Morphology and dissection of the two *Sthenoteuthis* species.** "Typical form" and "Dwarf form" of purpleback flying squids are referred to as *S. oualaniensis* and *Sthenoteuthis* sp., respectively. **A.** Morphology and the dorsal photophore patch of the "Typical form". **B.** Morphology and the corresponding position of the photophore (pseudo-photophore) of the "Dwarf form". **C.** Dissection of the female individual. **D.** Dissection of the male individual.

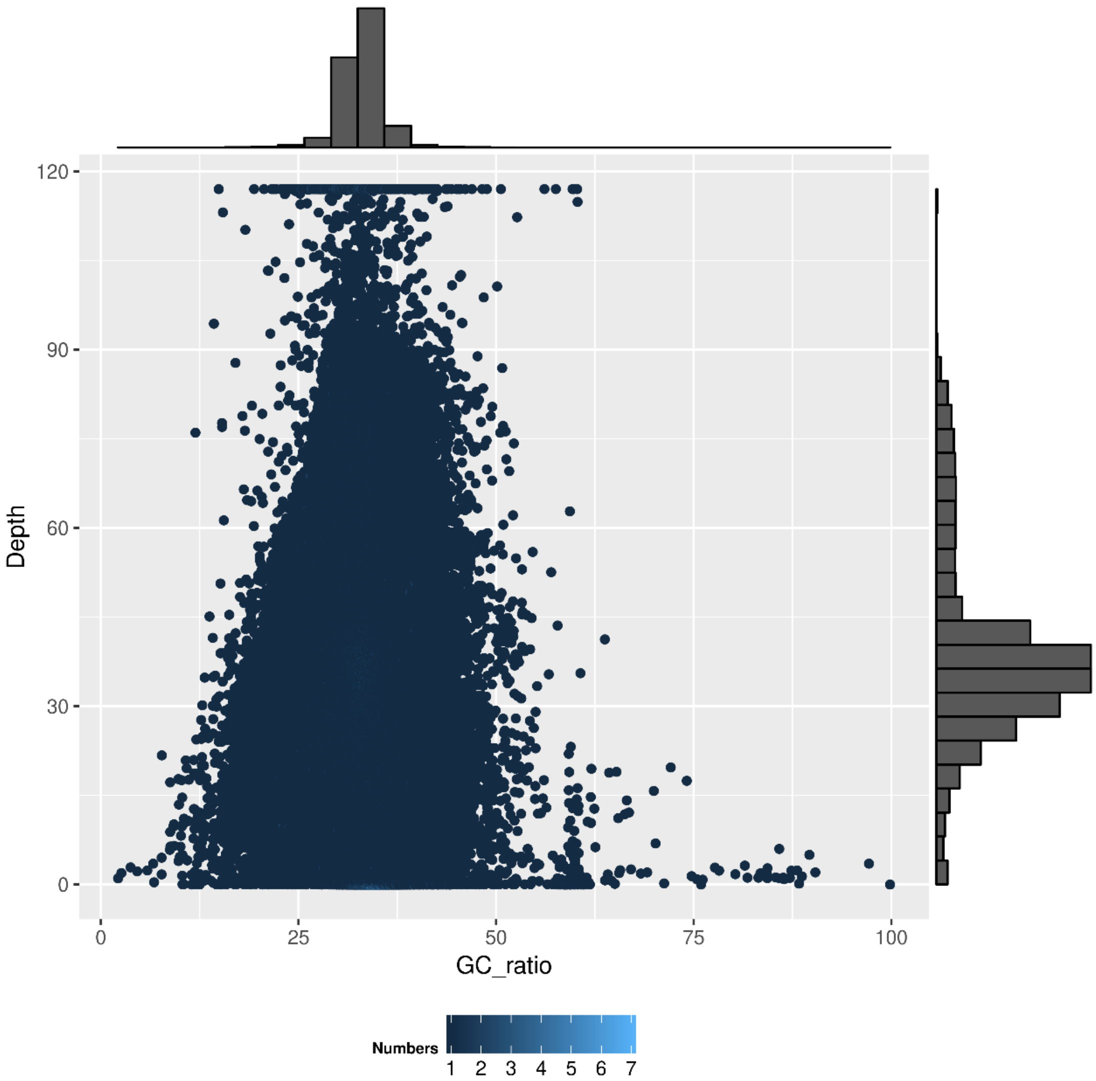


**Supplementary Figure S2 Distribution of GC contents versus the sequencing depth for *S. oualaniensis*.** The horizontal and vertical coordinates indicate the GC contents and sequencing depth respectively with 10 kb as a window. The GC contents of most windows were around 20%–50%, indicating the high-quality raw reads without significant contamination.

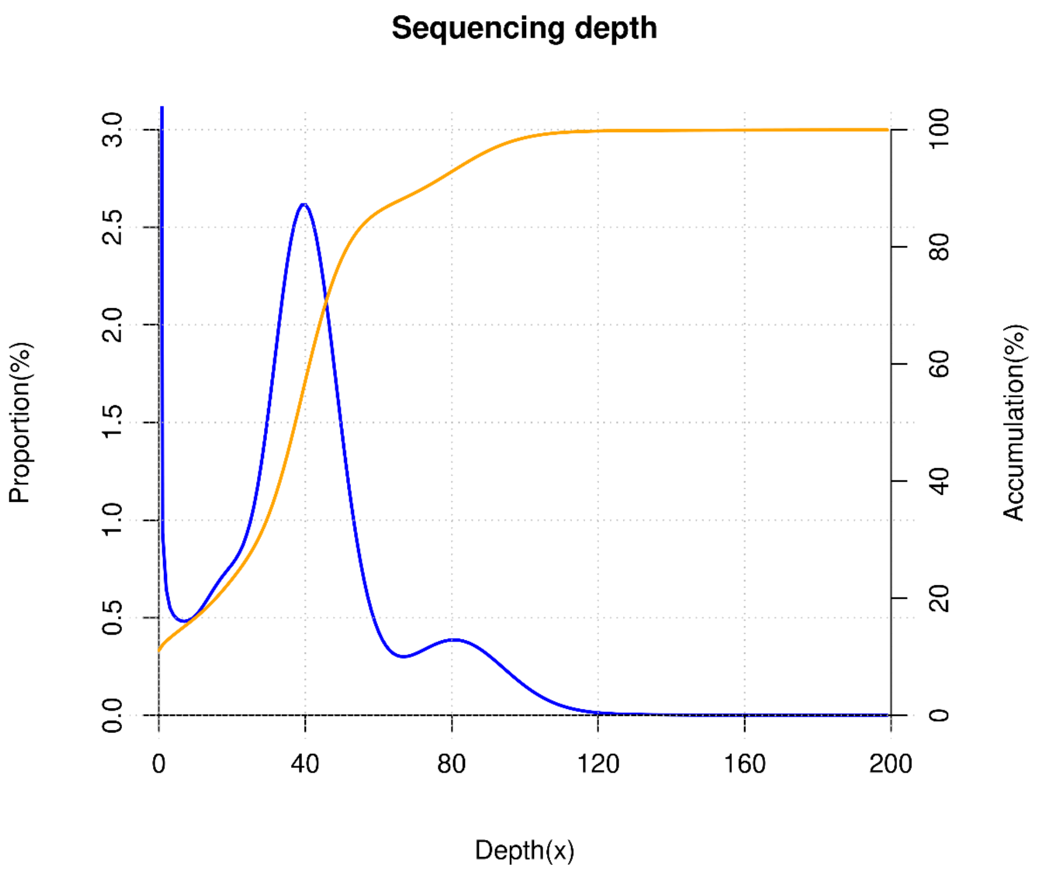


**Supplementary Figure S3 Sequencing depth distribution for *S. oualaniensis*.** The horizontal coordinate indicates the reads depth of PacBio reads. The vertical coordinate indicates the cumulative ratio of base depth. The blue line indicates the reads depth, the yellow line indicates the accumulation of all the reads depth.

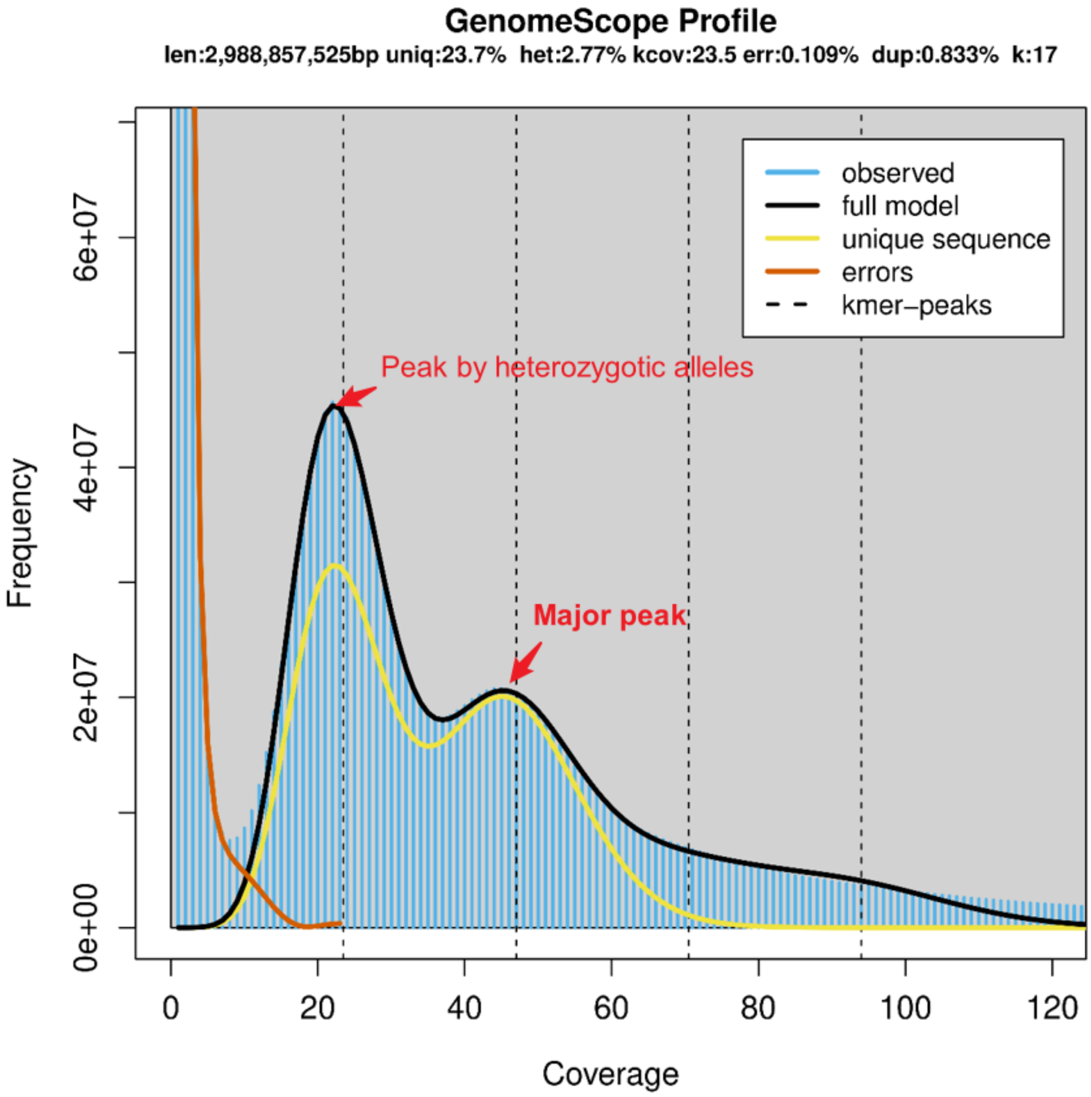


**Supplementary Figure S4 GenomeScope profiles for *S. oualaniensis* were estimated.** The result from jellyfish v2.2.6 with a *k*-mer size of 17 was used as the input file of GenomeScope v1.0. The second peak should be the major peak and has an estimated genome size of about 5.8 Gb. This is suggestive of the high heterozygosity of the species.

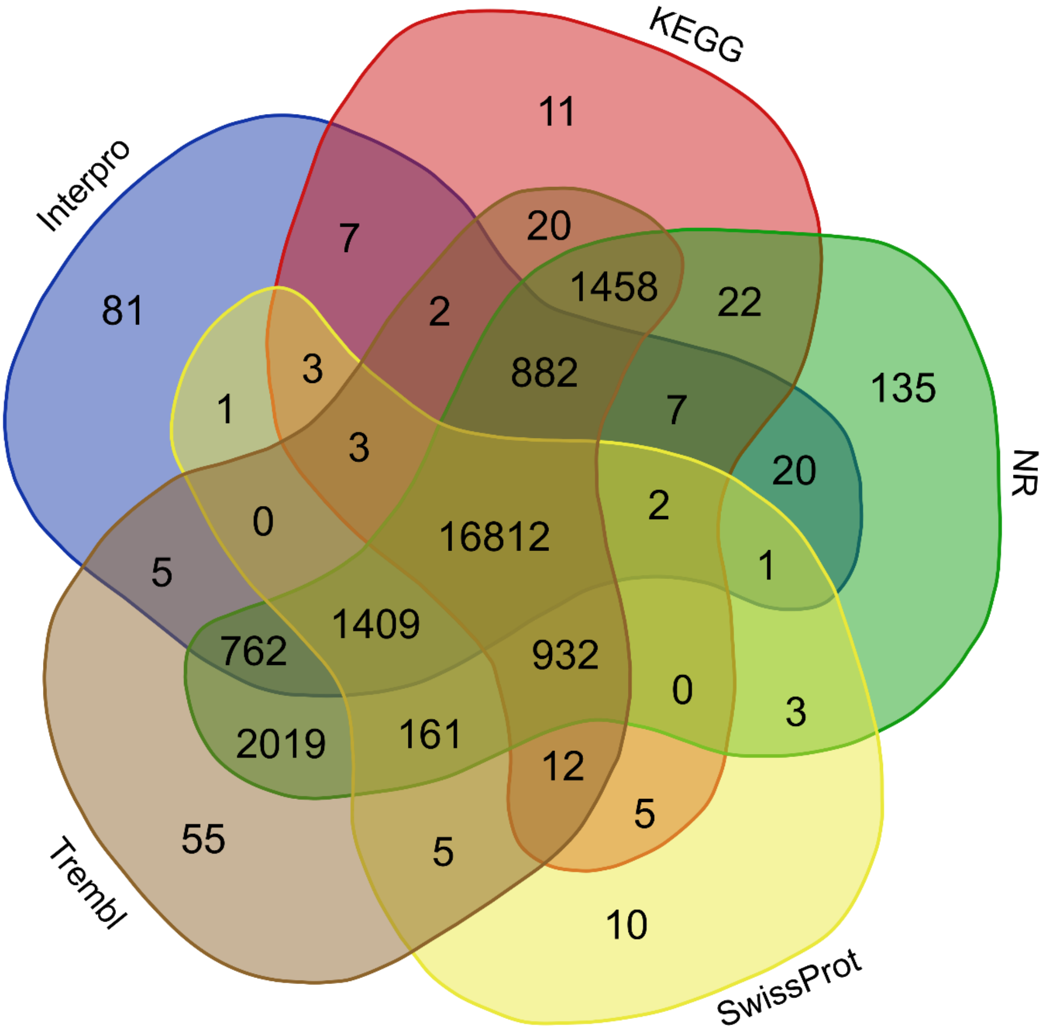


**Supplementary Figure S5** Venn diagram of *S. oualaniensis* to show the well-aligned gene sets from a different database, including NR, InterPro, KEGG, SwissProt, and Trembl database.

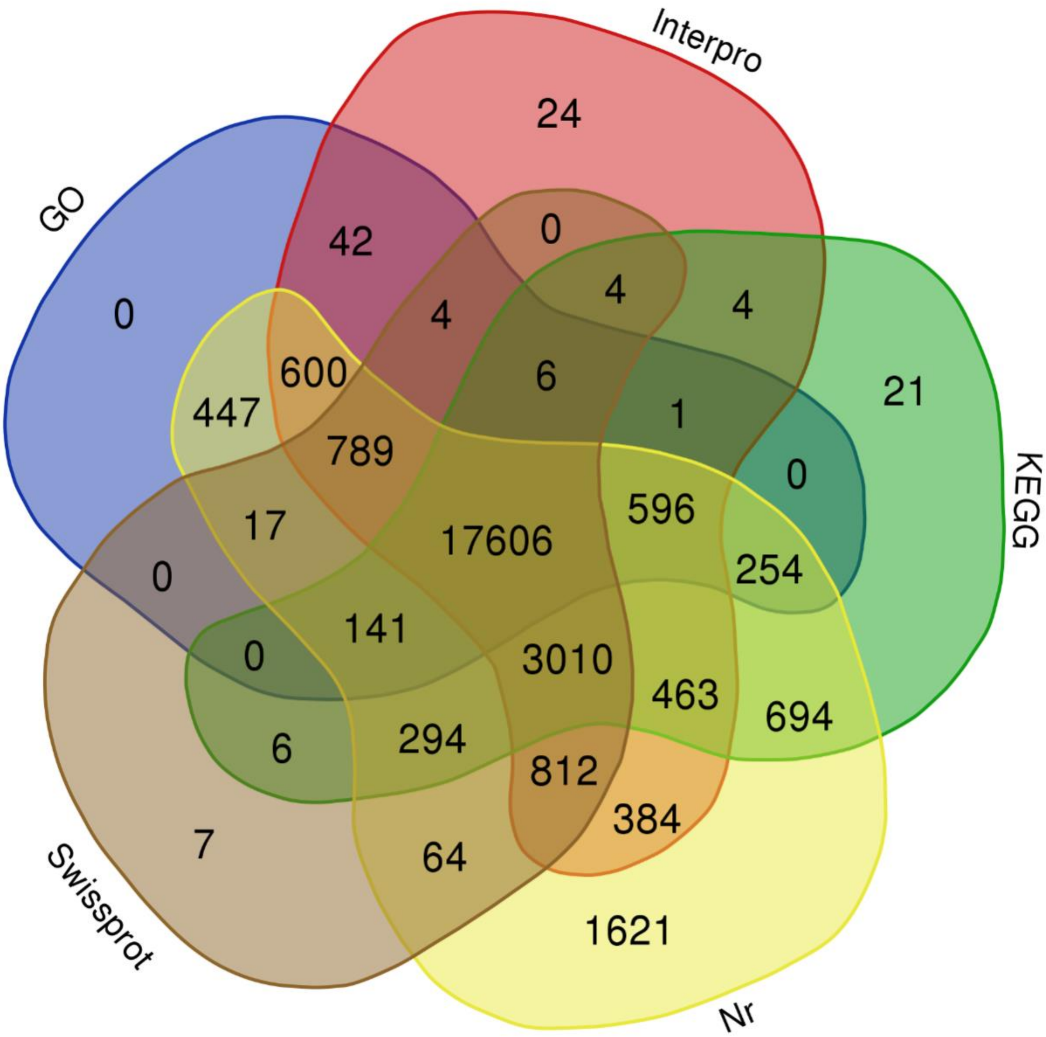


**Supplementary Figure S6** Venn diagram of *Sthenoteuthis* sp. to show the well-aligned gene sets from a different database, including NR, InterPro, KEGG, SwissProt, and Trembl database.

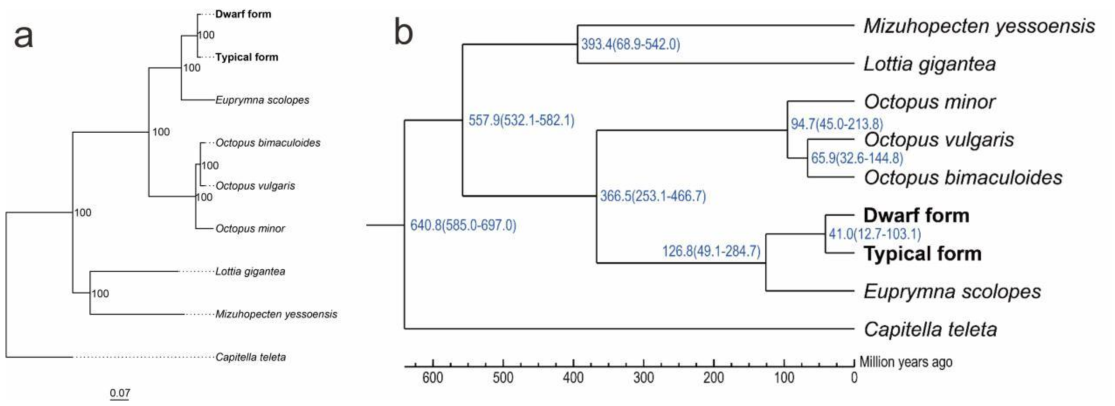


**Supplementary Figure S7** Estimation of **(A)** phylogeny and **(B)** divergence times among purpleback flying squid species and their relatives. The divergence time was indicated by the blue number near each node, while the 95% CI was indicated by the blue brackets and its specific value was colored in blue under the corresponding divergence time. Two softbound calibration time points had been applied: *Octopus bimaculoides* - *Capitella teleta* (585 - 679 MYA) and *Octopus bimaculoides* - *Lottia gigantea* (531 - 582 MYA).

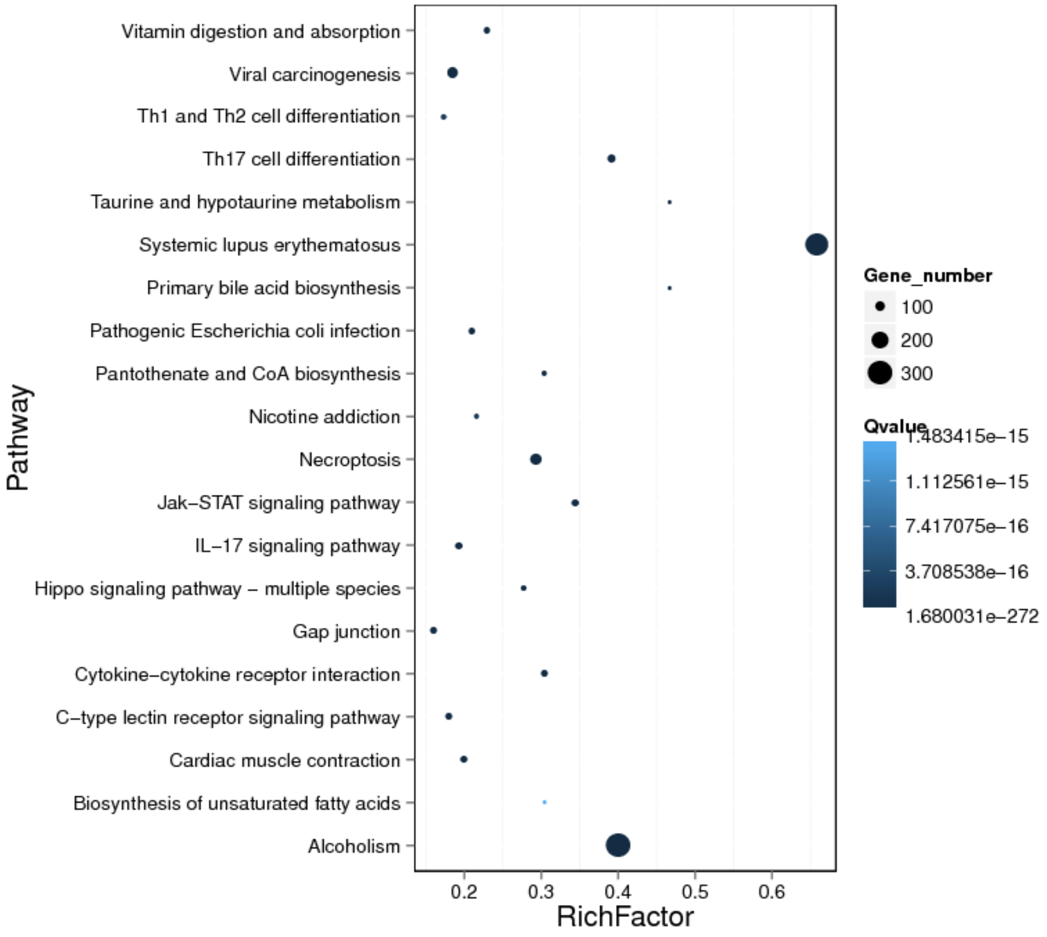


**Supplementary Figure S8** Top 20 KEGG enriched pathway of the significantly expanded gene families for both *Sthenoteuthis* species.

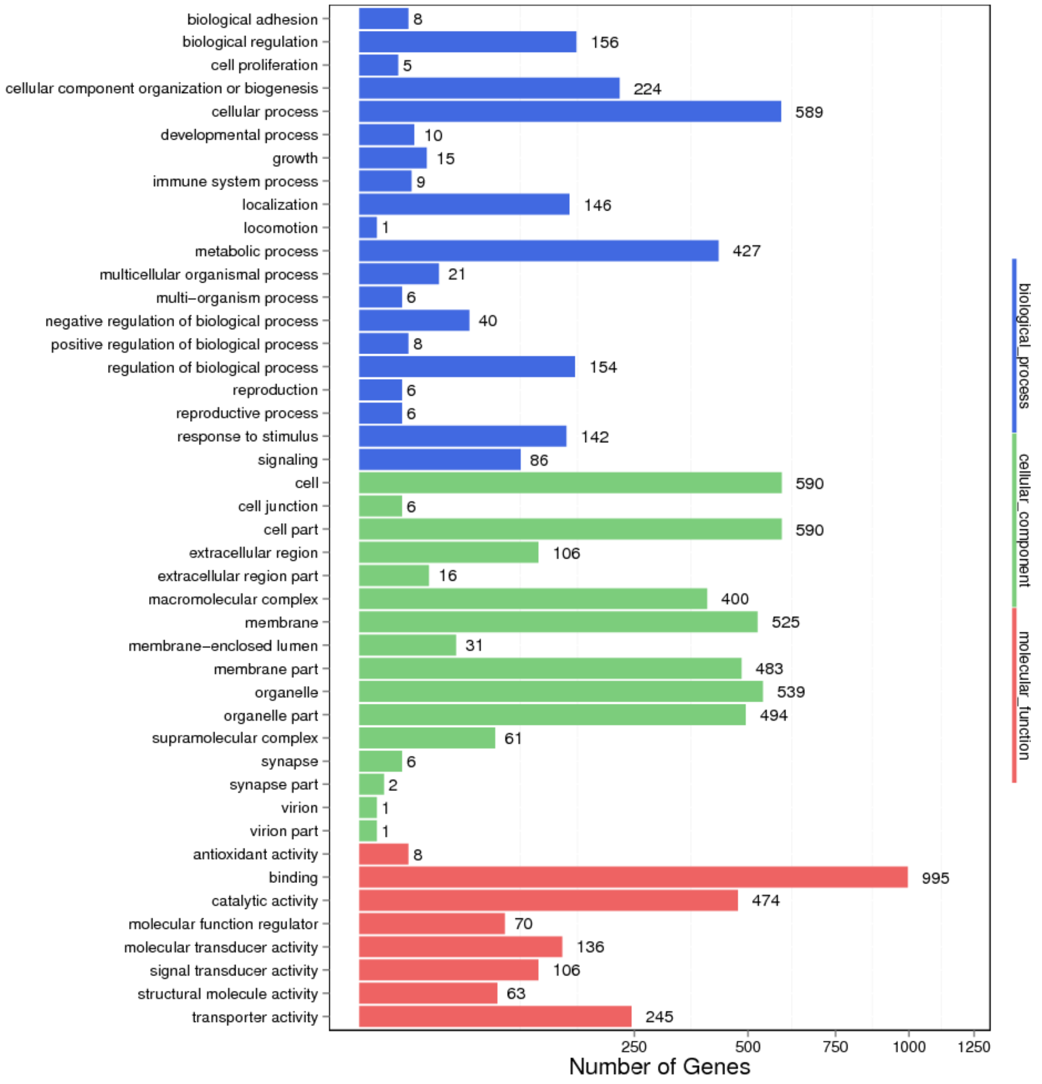


**Supplementary Figure S9** The significantly enriched GO terms of the expanded gene families for both *Sthenoteuthis* species.

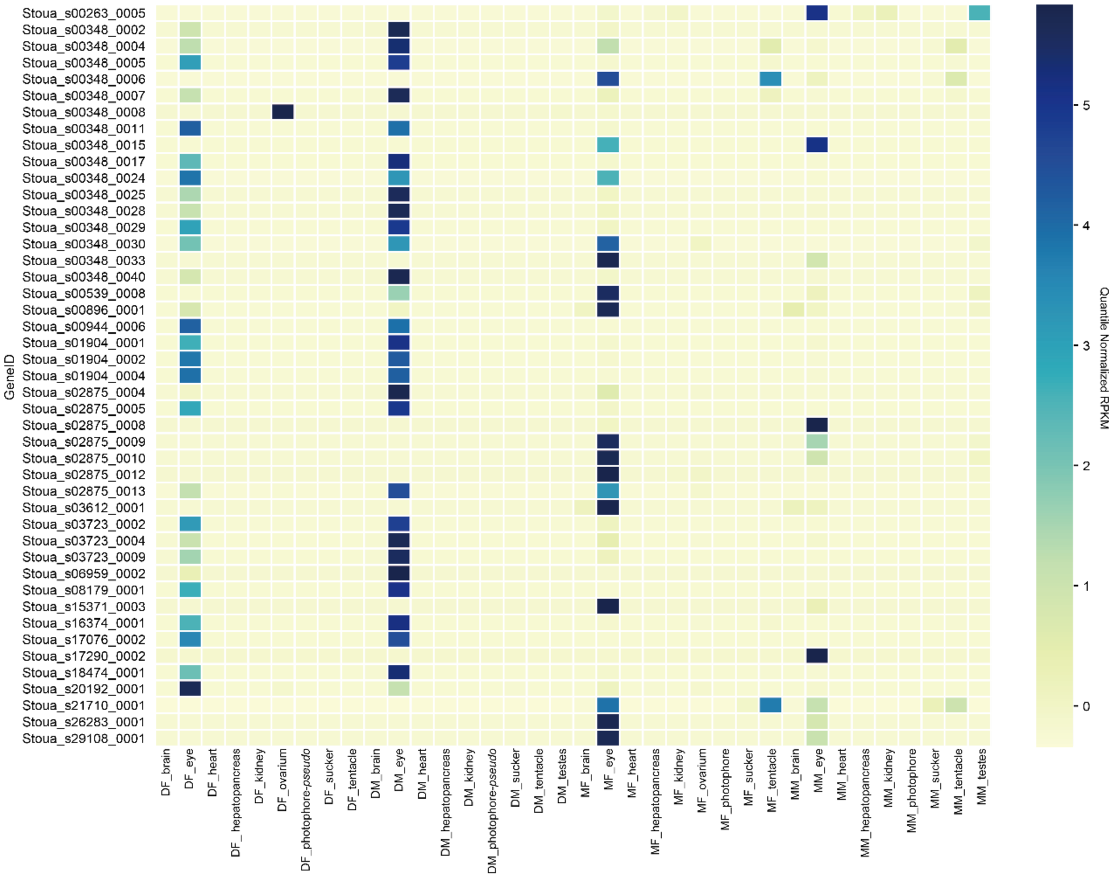


**Supplementary Figure S10 Expression patterns of coexisting gene members of the *SL20-1* subfamily in *S. oualaniensis* and *Sthenoteuthis* sp.** Only those expressed gene copies were shown. Gene IDs and tissue information correspond to Figure 3C. Most of the significantly expanded genes were highly expressed in the eyes.

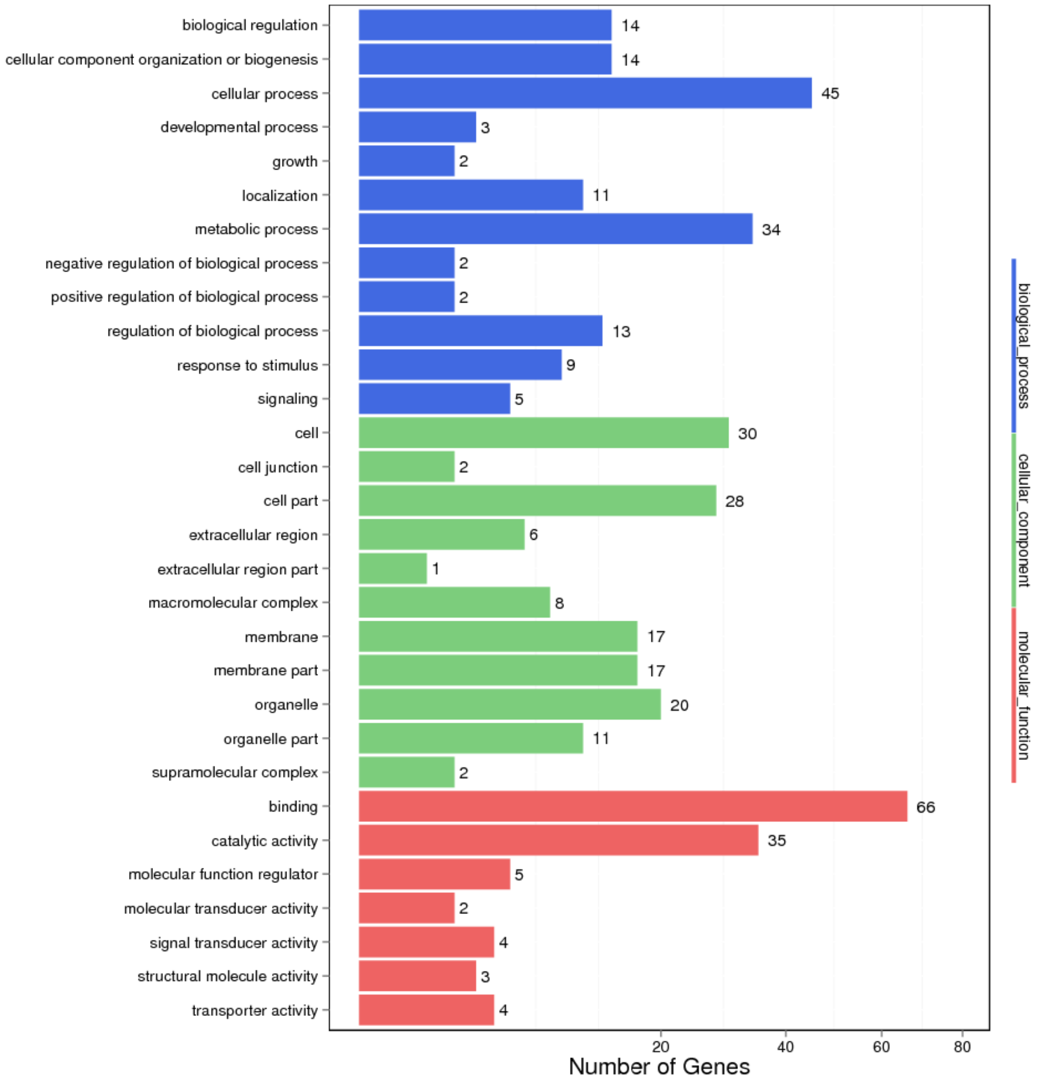


**Supplementary Figure S11** Classification of the three major GO function annotations for the 66 PSGs of the two *Stenoteuthis* lineage.

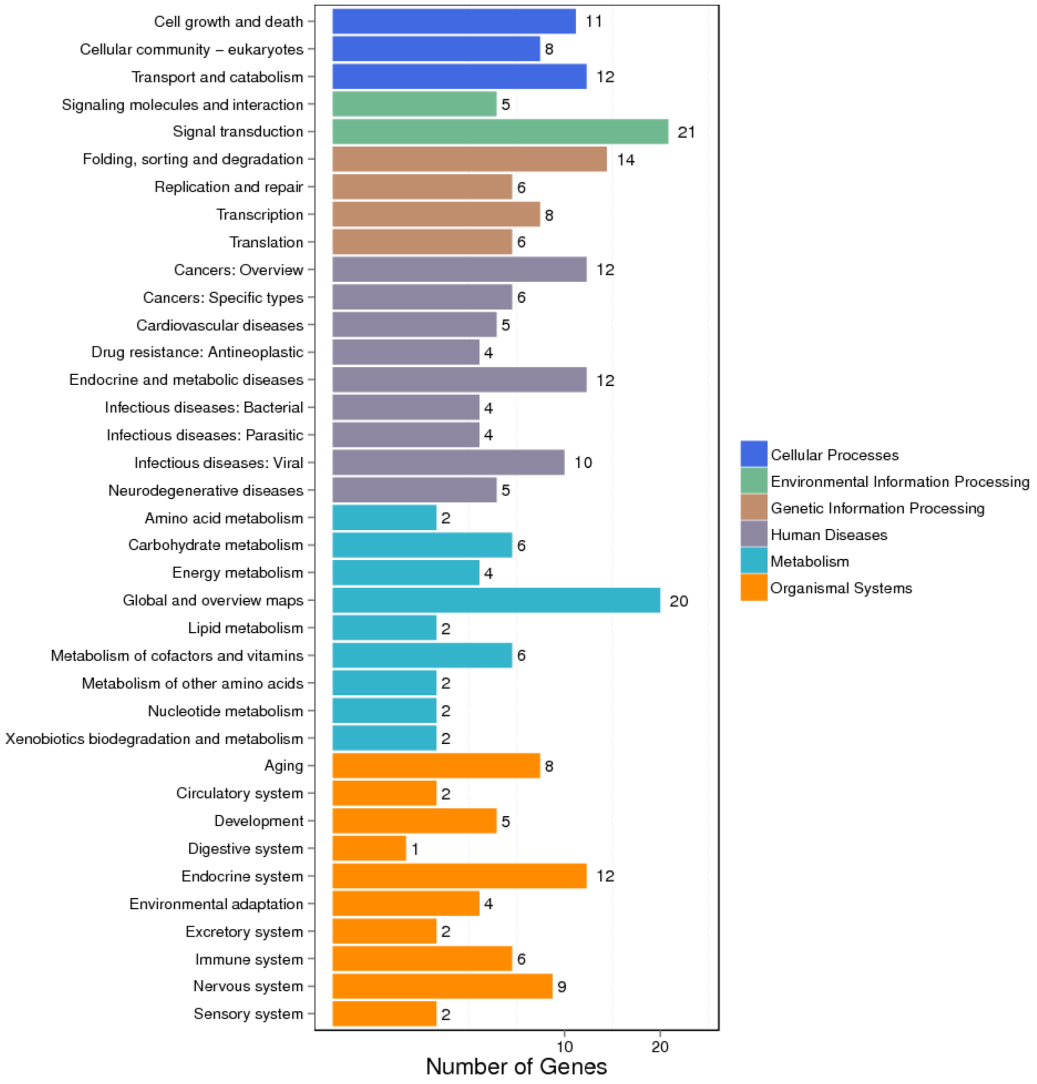


**Supplementary Figure S12** The significantly enriched KEGG pathway of the positively selected genes for both *Sthenoteuthis* species.

| Species/Abbrv | Group Name | * | * | | * | | | | * | | | | | | * | * | * | * | * | * | * | * | * | * | * | * | * |
|---|---|---|---|---|---|---|---|---|---|---|---|---|---|---|---|---|---|---|---|---|---|---|---|---|---|---|---|
| 1. Capitella_teleta | | R | V | N | D | A | M | M | P | Y | H | V | N | F | Y | G | N | P | H | S | R | T | H | A | Y | G | W |
| 2. Euprymna_scolopes | | R | V | L | D | A | M | L | P | Y | M | V | S | H | Y | G | N | P | H | S | R | T | H | A | Y | G | W |
| 3. Lottia_gigantea | | R | V | L | D | A | I | M | P | Y | Q | V | S | Y | Y | G | N | P | H | S | R | T | H | A | Y | G | W |
| 4. Mizuhopecten_yessoensis | | R | V | L | D | A | M | L | P | Y | F | V | S | Y | Y | G | N | P | H | S | R | T | H | A | Y | G | W |
| 5. Octopus_bimaculoides | | R | V | I | D | T | M | L | P | Y | M | I | S | F | Y | G | N | P | H | S | R | T | H | A | Y | G | W |
| 6. Octopus_minor | | R | V | L | D | A | M | L | P | Y | M | I | S | S | Y | G | N | P | H | S | R | T | H | A | Y | G | W |
| 7. Octopus_vulgaris | | R | V | I | D | T | M | L | P | Y | M | I | S | F | Y | G | N | P | H | S | R | T | H | A | Y | G | W |
| 8. Typical_form | | R | V | L | D | A | M | L | P | H | M | V | S | Y | Y | G | N | P | H | S | R | T | H | A | Y | G | W |
| 9. Dwarf_form | | R | V | L | D | A | M | L | P | H | M | V | S | Y | Y | G | N | P | H | S | R | T | H | A | Y | G | W |

**Supplementary Figure S13** Sequence alignments of *IscS* for the two *Sthenoteuthis* and other seven species from Mollusca and Annelida.

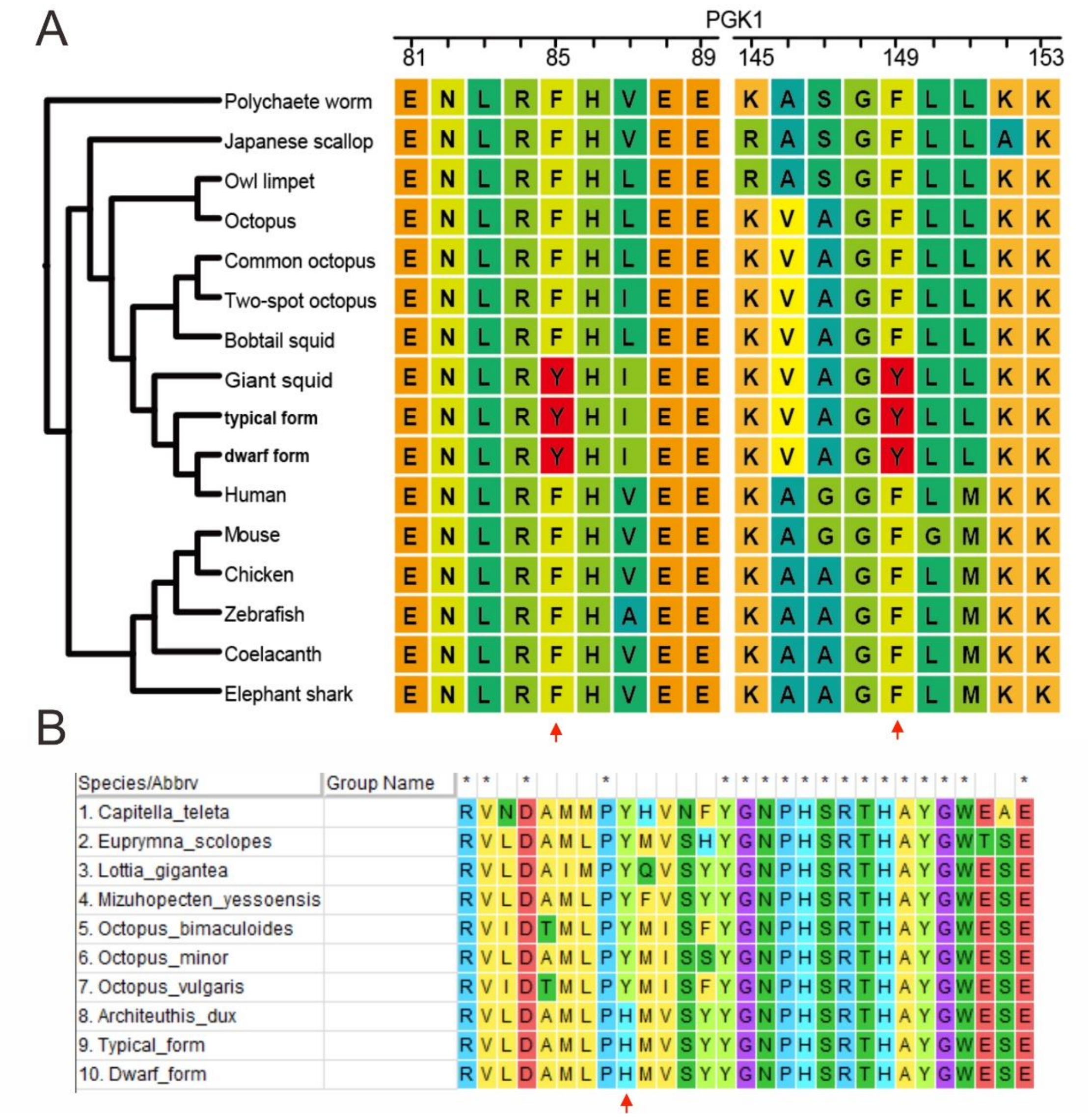


**Supplementary Figure S14** Positive selection sites for genes *PGK1* (Figure 4B) and *IscS* (Figure S13) shared by the two *Sthenoteuthis* were also detected from the giant squid (*Architeuthis dux*).

**Supplementary Table S1** PacBio reads statistics of the two *Sthenoteuthis* species.

| Library | Total Bases (Gb) | Total reads number | Average Length (bp) | Max Length (bp) | Min Length (bp) | N50 Length (bp) |
|---|---|---|---|---|---|---|
| r64048_20200304_014532-2_B01 | 196 | 14,755,426 | 13,305 | 263,237 | 50 | 16,488 |
| r64048_20200309_050911-4_D01 | 207 | 14,264,885 | 14,526 | 377,584 | 50 | 17,036 |

**Supplementary Table S2** Statistics of NGS sequencing.

| Platform | Lane name | Reads count | Base (bp) | Length | Q20 | Q30 | GC (%) |
|---|---|---|---|---|---|---|---|
| NGS | 171231_X602_FCH5GKMCCXY_L6_wHADPI064996-130 | 490,360,001 | 1.471E+11 | 150;150 | 96.72;93.78 | 92.33;86.93 | 34.63;34.91 |
| | 171231_X602_FCH5GKMCCXY_L7_wHADPI064996-130 | 490,187,979 | 1.471E+11 | 150;150 | 96.72;93.60 | 92.40;86.58 | 34.60;34.87 |
| | 171231_X602_FCH5GKMCCXY_L8_wHAIPI064995-133 | 490,049,960 | 1.47E+11 | 150;150 | 96.62;91.98 | 92.08;83.93 | 33.77;34.13 |
| | 171231_X602_FCH5GHTCCXY_L4_WHSTHwwxDEAADWAAPEI-32 | 484,213,479 | 1.453E+11 | 150;150 | 97.95;94.72 | 95.23;89.63 | 36.20;36.39 |
| | 171231_X602_FCH5GHTCCXY_L5_WHSTHwwxDEAADWAAPEI-32 | 484,008,671 | 1.452E+11 | 150;150 | 97.84;94.66 | 94.98;89.54 | 36.25;36.44 |
| | 171231_X602_FCH5GHTCCXY_L1_WHSTHwwxDFAADLAAPEI-92 | 469,606,815 | 1.409E+11 | 150;150 | 97.64;94.31 | 94.50;88.77 | 35.94;36.21 |
| | 171231_X602_FCH5GHTCCXY_L2_WHSTHwwxDFAADLAAPEI-92 | 468,872,994 | 1.407E+11 | 150;150 | 97.61;93.85 | 94.46;87.89 | 35.92;36.16 |
| | 171231_X602_FCH5GHTCCXY_L3_WHSTHwwxDGAADTAAPEI-84 | 480,129,452 | 1.44E+11 | 150;150 | 97.72;94.81 | 94.61;89.43 | 35.52;35.72 |
| | 171231_X602_FCH5GHTCCXY_L6_WHSTHwwxDHAADUAAPEI-86 | 480,589,371 | 1.442E+11 | 150;150 | 96.69;92.99 | 93.51;87.34 | 36.13;36.40 |
| | 180106_I188_FCCBT7CANXX_L2_wHAMPI064997-131 | 239,118,372 | 5.978E+10 | 125;125 | 92.85;90.13 | 86.31;82.17 | 33.35;33.50 |
| | 180106_I188_FCCBT7CANXX_L3_wHAMPI064997-131 | 239,565,426 | 5.989E+10 | 125;125 | 92.53;91.30 | 85.77;84.37 | 33.36;33.47 |
| | 180106_I188_FCCBT7CANXX_L4_wHAMPI064997-131 | 239,654,642 | 5.991E+10 | 125;125 | 93.08;91.29 | 86.76;84.39 | 33.36;33.48 |
| | 180106_I188_FCCBT7CANXX_L5_wHAMPI064997-131 | 240,438,866 | 6.011E+10 | 125;125 | 92.86;87.26 | 86.37;77.71 | 33.36;33.68 |
| | 171212_X603_FCH5GL3CCXY_L1_wHADPI064996-130 | 485,174,438 | 1.456E+11 | 150;150 | 97.83;96.22 | 94.67;90.86 | 34.37;34.52 |
| | 171226_X603_FCH5GL3CCXY_L1_wHADPI064996-130 | 497,996,920 | 1.494E+11 | 150;150 | 97.37;95.07 | 93.97;89.35 | 34.54;34.78 |
| | 171212_X603_FCH5GL3CCXY_L2_wHAIPI064995-133 | 477,806,525 | 1.433E+11 | 150;150 | 98.04;94.25 | 95.09;87.36 | 33.67;33.88 |
| | 171226_X603_FCH5GL3CCXY_L2_wHAIPI064995-133 | 511,818,498 | 1.535E+11 | 150;150 | 97.15;91.34 | 93.59;83.54 | 33.82;34.31 |
| | 171224_X603_FCH5GJ2CCXY_L6_WHSTHwwxDEAADWAAPEI-32 | 496,803,120 | 1.49E+11 | 150;150 | 97.76;94.21 | 94.83;88.54 | 36.26;36.39 |
| | 171224_X603_FCH5GJ2CCXY_L2_WHSTHwwxDFAADLAAPEI-92 | 490,644,651 | 1.472E+11 | 150;150 | 97.56;93.51 | 94.47;87.41 | 36.04;36.24 |
| | 171224_X603_FCH5GJ2CCXY_L3_WHSTHwwxDGAADTAAPEI-84 | 493,234,764 | 1.48E+11 | 150;150 | 97.51;94.07 | 94.30;87.99 | 35.61;35.74 |
| | 171224_X603_FCH5GJ2CCXY_L7_WHSTHwwxDHAADUAAPEI-86 | 490,555,492 | 1.472E+11 | 150;150 | 96.54;92.13 | 93.18;85.62 | 36.07;36.21 |
| | 171224_X603_FCH5GJ2CCXY_L4_WHSTHwwxDIAADVAAPEI-88 | 472,614,848 | 1.418E+11 | 150;150 | 96.66;92.94 | 93.15;86.79 | 34.83;35.18 |

**Supplementary Table S3** Statistics of 10x sequencing.

| Platform | Lane name | Reads count | Base (bp) | Length | Q20 | Q30 | GC |
|---|---|---|---|---|---|---|---|
| 10x | 180407_X495_FCHKG7KCCXY_L7_STHwwxD-AD-G10-1 | 41,845,197,900 | 4.18E+10 | 150;150 | 96.83;92.72 | 92.70;85.77 | 36.79;34.90 |
| | 180407_X495_FCHKG7KCCXY_L7_STHwwxD-AD-G10-2 | 29,492,050,800 | 2.95E+10 | 150;150 | 96.94;93.38 | 92.96;87.08 | 36.73;34.77 |
| | 180407_X495_FCHKG7KCCXY_L7_STHwwxD-AD-G10-3 | 25,025,630,400 | 2.5E+10 | 150;150 | 96.87;93.14 | 92.81;86.62 | 36.77;34.84 |
| | 180407_X495_FCHKG7KCCXY_L7_STHwwxD-AD-G10-4 | 29,713,203,600 | 2.97E+10 | 150;150 | 96.87;93.21 | 92.82;86.74 | 36.76;34.82 |
| | 180407_X495_FCHKGNNCCXY_L6_STHwwxD-AD-G10-1 | 42,188,685,300 | 4.22E+10 | 150;150 | 97.32;94.40 | 93.79;88.80 | 36.79;34.83 |
| | 180407_X495_FCHKGNNCCXY_L6_STHwwxD-AD-G10-2 | 29,634,741,900 | 2.96E+10 | 150;150 | 97.45;95.01 | 94.10;90.05 | 36.73;34.72 |
| | 180407_X495_FCHKGNNCCXY_L6_STHwwxD-AD-G10-3 | 25114150500 | 2.51E+10 | 150;150 | 97.35;94.78 | 93.89;89.61 | 36.76;34.77 |
| | 180407_X495_FCHKGNNCCXY_L6_STHwwxD-AD-G10-4 | 29,838,146,700 | 2.98E+10 | 150;150 | 97.37;94.86 | 93.91;89.73 | 36.75;34.75 |
| | 180407_X495_FCHKGNNCCXY_L7_STHwwxD-AD-G10-1 | 41,858,298,000 | 4.19E+10 | 150;150 | 97.35;94.27 | 93.85;88.57 | 36.79;34.85 |
| | 180407_X495_FCHKGNNCCXY_L7_STHwwxD-AD-G10-2 | 29,377,740,600 | 2.94E+10 | 150;150 | 97.48;94.90 | 94.15;89.84 | 36.72;34.73 |
| | 180407_X495_FCHKGNNCCXY_L7_STHwwxD-AD-G10-3 | 24,917,654,100 | 2.49E+10 | 150;150 | 97.38;94.67 | 93.94;89.40 | 36.76;34.78 |
| | 180407_X495_FCHKGNNCCXY_L7_STHwwxD-AD-G10-4 | 29,626,963,800 | 2.96E+10 | 150;150 | 97.39;94.74 | 93.97;89.52 | 36.75;34.77 |
| | 180407_X528_FCHKGMVCCXY_L3_STHwwxD-AD-G10-1 | 42,281,920,200 | 4.23E+10 | 150;150 | 97.10;93.67 | 93.35;87.83 | 36.84;34.95 |
| | 180407_X528_FCHKGMVCCXY_L3_STHwwxD-AD-G10-2 | 29,725,274,100 | 2.97E+10 | 150;150 | 97.21;94.21 | 93.61;88.91 | 36.78;34.83 |
| | 180407_X528_FCHKGMVCCXY_L3_STHwwxD-AD-G10-3 | 25165895400 | 2.52E+10 | 150;150 | 97.13;93.99 | 93.45;88.49 | 36.81;34.90 |
| | 180407_X528_FCHKGMVCCXY_L3_STHwwxD-AD-G10-4 | 29,904,066,600 | 2.99E+10 | 150;150 | 97.14;94.06 | 93.47;88.60 | 36.80;34.88 |
| | 180407_X528_FCHKGMVCCXY_L4_STHwwxD-AD-G10-1 | 42,916,878,600 | 4.29E+10 | 150;150 | 97.21;93.91 | 93.61;88.35 | 36.84;34.96 |
| | 180407_X528_FCHKGMVCCXY_L4_STHwwxD-AD-G10-2 | 30,144,276,300 | 3.01E+10 | 150;150 | 97.31;94.43 | 93.86;89.39 | 36.78;34.84 |
| | 180407_X528_FCHKGMVCCXY_L4_STHwwxD-AD-G10-3 | 25,489,609,800 | 2.55E+10 | 150;150 | 97.23;94.22 | 93.70;88.99 | 36.82;34.91 |
| | 180407_X528_FCHKGMVCCXY_L4_STHwwxD-AD-G10-4 | 30305369700 | 3.03E+10 | 150;150 | 97.25;94.30 | 93.73;89.12 | 36.81;34.88 |
| | 180407_X528_FCHKGMVCCXY_L5_STHwwxD-AD-G10-1 | 43,003,608,900 | 4.3E+10 | 150;150 | 97.23;94.01 | 93.67;88.48 | 36.83;34.94 |
| | 180407_X528_FCHKGMVCCXY_L5_STHwwxD-AD-G10-2 | 30,197,941,200 | 3.02E+10 | 150;150 | 97.33;94.54 | 93.91;89.54 | 36.77;34.82 |
| | 180407_X528_FCHKGMVCCXY_L5_STHwwxD-AD-G10-3 | 25,552,044,900 | 2.56E+10 | 150;150 | 97.25;94.33 | 93.75;89.15 | 36.81;34.88 |
| | 180407_X528_FCHKGMVCCXY_L5_STHwwxD-AD-G10-4 | 30,391,401,000 | 3.04E+10 | 150;150 | 97.27;94.41 | 93.78;89.29 | 36.80;34.86 |

**Supplementary Table S4** Statistics of Hi-C sequencing.

| Platform | Lane name | reads | Base | Length | Q20 | Q30 | GC |
|---|---|---|---|---|---|---|---|
| | 190323_I25_V300014809_L1_WHRDSTHwwxDAAAMAB-509 | 51,681,097 | 15,504,329,100 | 150;150 | 96.23;94.65 | 87.93;84.58 | 35.86;36.01 |
| | 190323_I25_V300014809_L1_WHRDSTHwwxDAAAMAB-510 | 60,349,538 | 18,104,861,400 | 150;150 | 96.14;94.42 | 87.71;84.16 | 35.87;36.01 |
| | 190323_I25_V300014809_L1_WHRDSTHwwxDAAAMAB-511 | 64,110,433 | 19,233,129,900 | 150;150 | 96.23;94.89 | 87.98;85.19 | 35.86;35.99 |
| | 190323_I25_V300014809_L1_WHRDSTHwwxDAAAMAB-512 | 59,664,912 | 17,899,473,600 | 150;150 | 96.32;95.24 | 88.24;86.00 | 35.87;35.96 |
| | 190323_I25_V300014809_L1_WHRDSTHwwxDAAAMAB-513 | 47,233,033 | 14,169,909,900 | 150;150 | 96.30;94.97 | 88.19;85.47 | 35.82;35.96 |
| | 190323_I25_V300014809_L1_WHRDSTHwwxDAAAMAB-514 | 54,825,110 | 16,447,533,000 | 150;150 | 96.05;95.06 | 87.56;85.56 | 35.89;35.98 |
| | 190323_I25_V300014809_L1_WHRDSTHwwxDAAAMAB-515 | 52,303,566 | 15,691,069,800 | 150;150 | 96.19;94.69 | 87.83;84.74 | 35.86;36.02 |
| | 190323_I25_V300014809_L1_WHRDSTHwwxDAAAMAB-516 | 66,507,454 | 19,952,236,200 | 150;150 | 96.17;94.68 | 87.80;84.70 | 35.91;36.02 |
| | 190113_I412_CL100111910_L1_WHRDSTHwwxDAAAMAB-509 | 62,028,564 | 12,405,712,800 | 100;100 | 97.49;92.99 | 90.16;80.48 | 35.57;35.56 |
| | 190113_I412_CL100111910_L1_WHRDSTHwwxDAAAMAB-510 | 74,317,337 | 14,863,467,400 | 100;100 | 97.53;93.17 | 90.26;80.87 | 35.59;35.52 |
| | 190113_I412_CL100111910_L1_WHRDSTHwwxDAAAMAB-511 | 78071192 | 15,614,238,400 | 100;100 | 97.55;93.33 | 90.32;81.24 | 35.58;35.54 |
| | 190113_I412_CL100111910_L1_WHRDSTHwwxDAAAMAB-512 | 74,175,906 | 14,835,181,200 | 100;100 | 97.58;93.34 | 90.43;81.26 | 35.58;35.53 |
| | 190113_I412_CL100111910_L1_WHRDSTHwwxDAAAMAB-513 | 57,232,743 | 11,446,548,600 | 100;100 | 97.52;93.17 | 90.24;80.93 | 35.54;35.54 |
| | 190113_I412_CL100111910_L1_WHRDSTHwwxDAAAMAB-514 | 67,704,437 | 13,540,887,400 | 100;100 | 97.40;93.16 | 89.93;80.89 | 35.60;35.56 |
| | 190113_I412_CL100111910_L1_WHRDSTHwwxDAAAMAB-515 | 64,249,139 | 12,849,827,800 | 100;100 | 97.57;93.45 | 90.38;81.52 | 35.57;35.55 |
| | 190113_I412_CL100111910_L1_WHRDSTHwwxDAAAMAB-516 | 81,973,766 | 16,394,753,200 | 100;100 | 97.45;93.25 | 90.06;81.05 | 35.62;35.53 |
| | 190312_I525_CL100110998_L1_WHRDSTHwwxDAAAMAB-509 | 73,023,423 | 14,604,684,600 | 100;100 | 96.76;91.59 | 88.76;77.58 | 35.63;35.49 |
| | 190312_I525_CL100110998_L1_WHRDSTHwwxDAAAMAB-510 | 87,849,948 | 17,569,989,600 | 100;100 | 96.62;91.60 | 88.40;77.65 | 35.68;35.50 |
| | 190312_I525_CL100110998_L1_WHRDSTHwwxDAAAMAB-511 | 94,088,826 | 18,817,765,200 | 100;100 | 96.85;91.98 | 89.02;78.44 | 35.63;35.49 |
| | 190312_I525_CL100110998_L1_WHRDSTHwwxDAAAMAB-512 | 89,867,469 | 17,973,493,800 | 100;100 | 96.82;92.15 | 88.94;78.78 | 35.66;35.50 |
| Hi-C | 190312_I525_CL100110998_L1_WHRDSTHwwxDAAAMAB-513 | 68,964,948 | 13,792,989,600 | 100;100 | 96.86;91.66 | 89.12;77.83 | 35.59;35.47 |
| | 190312_I525_CL100110998_L1_WHRDSTHwwxDAAAMAB-514 | 78,554,916 | 15,710,983,200 | 100;100 | 95.70;91.63 | 86.04;77.68 | 35.74;35.52 |
| | 190312_I525_CL100110998_L1_WHRDSTHwwxDAAAMAB-515 | 77,209,126 | 15,441,825,200 | 100;100 | 96.84;92.09 | 88.97;78.69 | 35.64;35.53 |
| | 190312_I525_CL100110998_L1_WHRDSTHwwxDAAAMAB-516 | 97,383,520 | 19,476,704,000 | 100;100 | 96.21;92.07 | 87.31;78.58 | 35.73;35.49 |
| | 190312_I525_CL100110998_L2_WHRDSTHwwxDAAAMAB-509 | 79,137,857 | 15,827,571,400 | 100;100 | 97.31;92.37 | 89.56;79.06 | 35.58;35.50 |
| | 190312_I525_CL100110998_L2_WHRDSTHwwxDAAAMAB-510 | 93,097,243 | 18,619,448,600 | 100;100 | 97.27;92.43 | 89.46;79.23 | 35.62;35.48 |
| | 190312_I525_CL100110998_L2_WHRDSTHwwxDAAAMAB-511 | 101,327,211 | 20,265,442,200 | 100;100 | 97.40;92.73 | 89.86;79.91 | 35.59;35.48 |
| | 190312_I525_CL100110998_L2_WHRDSTHwwxDAAAMAB-512 | 95,234,920 | 19,046,984,000 | 100;100 | 97.39;92.80 | 89.81;80.03 | 35.61;35.49 |
| | 190312_I525_CL100110998_L2_WHRDSTHwwxDAAAMAB-513 | 73,296,020 | 14,659,204,000 | 100;100 | 97.37;92.40 | 89.75;79.16 | 35.55;35.48 |
| | 190312_I525_CL100110998_L2_WHRDSTHwwxDAAAMAB-514 | 84,710,041 | 16,942,008,200 | 100;100 | 96.68;92.65 | 87.90;79.69 | 35.64;35.51 |
| | 190312_I525_CL100110998_L2_WHRDSTHwwxDAAAMAB-515 | 82,677,852 | 16,535,570,400 | 100;100 | 97.40;92.86 | 89.85;80.20 | 35.59;35.50 |
| | 190312_I525_CL100110998_L2_WHRDSTHwwxDAAAMAB-516 | 104,695,019 | 20,939,003,800 | 100;100 | 96.99;92.82 | 88.72;80.06 | 35.66;35.48 |
| | 190312_I569_CL100112082_L1_WHRDSTHwwxDAAAMAB-509 | 75,811,112 | 15,162,222,400 | 100;100 | 97.53;93.13 | 89.76;80.68 | 35.58;35.56 |
| | 190312_I569_CL100112082_L1_WHRDSTHwwxDAAAMAB-510 | 89,608,806 | 17,921,761,200 | 100;100 | 97.51;93.14 | 89.70;80.78 | 35.61;35.54 |
| | 190312_I569_CL100112082_L1_WHRDSTHwwxDAAAMAB-511 | 96,092,960 | 19,218,592,000 | 100;100 | 97.57;93.43 | 89.90;81.38 | 35.60;35.55 |
| | 190312_I569_CL100112082_L1_WHRDSTHwwxDAAAMAB-512 | 90,603,751 | 18,120,750,200 | 100;100 | 97.59;93.49 | 89.95;81.49 | 35.61;35.55 |
| | 190312_I569_CL100112082_L1_WHRDSTHwwxDAAAMAB-513 | 70,591,166 | 14,118,233,200 | 100;100 | 97.53;93.20 | 89.78;80.93 | 35.55;35.55 |
| | 190312_I569_CL100112082_L1_WHRDSTHwwxDAAAMAB-514 | 81,589,261 | 16,317,852,200 | 100;100 | 97.38;93.21 | 89.36;80.89 | 35.62;35.58 |
| | 190312_I569_CL100112082_L1_WHRDSTHwwxDAAAMAB-515 | 78,397,566 | 15,679,513,200 | 100;100 | 97.59;93.54 | 89.96;81.63 | 35.59;35.57 |
| | 190312_I569_CL100112082_L1_WHRDSTHwwxDAAAMAB-516 | 99,394,009 | 19,878,801,800 | 100;100 | 97.42;93.44 | 89.47;81.39 | 35.65;35.55 |
| | 190312_I569_CL100112082_L2_WHRDSTHwwxDAAAMAB-509 | 77,644,100 | 15,528,820,000 | 100;100 | 97.49;93.36 | 89.83;80.86 | 35.60;35.53 |

| | | | | | | |
|---|---|---|---|---|---|---|
| 190312_I569_CL100112082_L2_WHRDSTHwwxDAAAMAB-510 | 90,986,991 | 18,197,398,200 | 100;100 | 97.49;93.39 | 89.83;80.98 | 35.63;35.51 |
| 190312_I569_CL100112082_L2_WHRDSTHwwxDAAAMAB-511 | 97,773,990 | 19,554,798,000 | 100;100 | 97.53;93.64 | 89.97;81.54 | 35.62;35.52 |
| 190312_I569_CL100112082_L2_WHRDSTHwwxDAAAMAB-512 | 91,650,661 | 18,330,132,200 | 100;100 | 97.55;93.69 | 90.01;81.62 | 35.63;35.52 |
| 190312_I569_CL100112082_L2_WHRDSTHwwxDAAAMAB-513 | 72,167,629 | 14,433,525,800 | 100;100 | 97.50;93.47 | 89.87;81.16 | 35.57;35.52 |
| 190312_I569_CL100112082_L2_WHRDSTHwwxDAAAMAB-514 | 82,894,530 | 16,578,906,000 | 100;100 | 97.35;93.44 | 89.45;81.08 | 35.64;35.55 |
| 190312_I569_CL100112082_L2_WHRDSTHwwxDAAAMAB-515 | 79,691,338 | 15,938,267,600 | 100;100 | 97.56;93.74 | 90.05;81.78 | 35.61;35.54 |
| 190312_I569_CL100112082_L2_WHRDSTHwwxDAAAMAB-516 | 101,040,813 | 20,208,162,600 | 100;100 | 97.40;93.64 | 89.59;81.51 | 35.67;35.53 |
| 190313_I101_CL100110996_L1_WHRDSTHwwxDAAAMAB-509 | 71,973,464 | 14,394,692,800 | 100;100 | 96.87;92.91 | 88.22;79.14 | 35.59;35.51 |
| 190313_I101_CL100110996_L1_WHRDSTHwwxDAAAMAB-510 | 84,325,203 | 16,865,040,600 | 100;100 | 96.82;92.76 | 88.07;78.92 | 35.61;35.50 |
| 190313_I101_CL100110996_L1_WHRDSTHwwxDAAAMAB-511 | 91,855,713 | 18,371,142,600 | 100;100 | 96.90;93.18 | 88.37;79.82 | 35.60;35.50 |
| 190313_I101_CL100110996_L1_WHRDSTHwwxDAAAMAB-512 | 85,194,550 | 17,038,910,000 | 100;100 | 96.96;93.34 | 88.50;80.08 | 35.61;35.51 |
| 190313_I101_CL100110996_L1_WHRDSTHwwxDAAAMAB-513 | 65,864,264 | 13,172,852,800 | 100;100 | 96.86;92.80 | 88.26;79.01 | 35.56;35.50 |
| 190313_I101_CL100110996_L1_WHRDSTHwwxDAAAMAB-514 | 77,782,321 | 15,556,464,200 | 100;100 | 96.76;92.99 | 87.98;79.37 | 35.63;35.54 |
| 190313_I101_CL100110996_L1_WHRDSTHwwxDAAAMAB-515 | 75,250,064 | 15,050,012,800 | 100;100 | 96.92;93.28 | 88.39;80.09 | 35.59;35.51 |
| 190313_I101_CL100110996_L1_WHRDSTHwwxDAAAMAB-516 | 96,708,875 | 19,341,775,000 | 100;100 | 96.79;93.19 | 88.02;79.84 | 35.65;35.50 |

**Supplementary Table S5** Statistics of the predicted gene sets for two *Sthenoteuthis* species and the comparison with other relatives.

| Species | Average mRNA Length | Average CDS Length | average Exon Length | Average Exon Number | Average Intron Length | Gene number | GC content |
|---|---|---|---|---|---|---|---|
| *Octopus bimaculoides* | 16,136.23 | 906.34 | 207.69 | 4.36 | 3,840.51 | 26,936 | 29.90% |
| *Euprymna scolopes* | 18,007.25 | 864.75 | 232.14 | 3.73 | 6,290.36 | 33,793 | 21.80% |
| *Octopus vulgaris* | 71,229.43 | 1,815.68 | 186.36 | 9.74 | 6,569.61 | 25,643 | 29.90% |
| *Sthenoteuthis oualaniensis* | 26,772.23 | 1,174.35 | 201.84 | 5.82 | 4,603.80 | 26,646 | 33.07% |
| *Sthenoteuthis sp.* | 36,141.98 | 1,250.93 | 180.68 | 6.92 | 4,878.78 | 28,715 | 33.00% |

**Supplementary Table S6** The sequencing reads information used for transcriptomes assembly from nine tissues.

| species | Sample | Reads counts | Average length | Min length | Max length |
|---|---|---|---|---|---|
| *S. oualaniensis* (Typical form) | Female ovary and male testes | 45,177 | 1324.66 | 200 | 25,632 |
| | Female and male brain | 56,385 | 1347.76 | 200 | 23,014 |
| | Female and male eye | 41,805 | 1493.96 | 200 | 23,013 |
| | Female and male heart | 28,352 | 1475.56 | 200 | 18,503 |
| | Female and male hepatopancreas | 35,243 | 1275.35 | 200 | 22,594 |
| | Female and male kidney | 36,832 | 1413.95 | 200 | 22,606 |
| | Female and male photophore | 27,673 | 1427.77 | 200 | 22,571 |
| | Female and male sucker | 43,903 | 1438.38 | 200 | 22,648 |
| | Female and male tentacle | 36,010 | 1455.85 | 200 | 22,588 |
| | Female and male testes | 38,574 | 1266.29 | 200 | 24,622 |
| *Sthenoteuthis sp.* (Dwarf form) | Female ovary and male testes | 48,255 | 1476.44 | 200 | 20,844 |
| | Female and male brain | 56,150 | 1549.72 | 200 | 22,569 |
| | Female and male eye | 40,463 | 1595.88 | 200 | 20,872 |
| | Female and male heart | 31,856 | 1547.5 | 200 | 24,737 |
| | Female and male hepatopancreas | 39,689 | 1363.96 | 200 | 19,782 |
| | Female and male kidney | 40,881 | 1563.33 | 200 | 31,105 |
| | Female and male photophore | 26,888 | 1512.01 | 200 | 23,903 |
| | Female and male sucker | 48,632 | 1531.42 | 200 | 16,809 |
| | Female and male tentacle | 40,475 | 1553.02 | 200 | 26,308 |
| | Female and male testes | 37,075 | 1447.34 | 200 | 25,850 |

**Supplementary Table S7** The predicted genome completeness of the two *Sthenoteuthis* species with metazoa_odb10.

| | ***S. oualaniensis*** | | ***Sthenoteuthis* sp.** | |
|---|---|---|---|---|
| | Number | Percentage (%) | Number | Percentage (%) |
| Total BUSCO groups searched | 954 | 100 | 954 | 100 |
| Complete BUSCOs (C) | 853 | 89.4 | 893 | 93.6 |
| Complete and single-copy BUSCOs (S) | 580 | 60.8 | 441 | 46.2 |
| Complete and duplicated BUSCOs (D) | 273 | 28.6 | 452 | 47.4 |
| Fragmented BUSCOs (F) | 28 | 2.9 | 19 | 2.0 |
| Missing BUSCOs (M) | 73 | 7.7 | 42 | 4.4 |

**Supplementary Table S8** Transposable elements information for the genome of *S. oualaniensis*.

| | Repbase TEs | | TE proteins | | De novo | | Combined TEs | |
|---|---|---|---|---|---|---|---|---|
| Type | Length (Bp) | Percent in genome | Length (Bp) | Percent in genome | Length (Bp) | Percent in genome | Length (Bp) | Percent in genome |
| DNA | 558,511,598 | 9.81 | 12,494,633 | 0.22 | 968,594,716 | 17.01 | 1,450,568,773 | 25.47 |
| LINE | 393,736,244 | 6.91 | 25,155,9979 | 4.42 | 1,119,402,772 | 19.66 | 1,255,803,973 | 22.05 |
| SINE | 13,618,899 | 0.24 | 0 | 0 | 56,359,932 | 0.99 | 69,532,719 | 1.22 |
| LTR | 15,015,9521 | 2.64 | 30,903,245 | 0.54 | 320,972,901 | 5.64 | 446,387,651 | 7.84 |
| Other | 908,764 | 0.02 | 0 | 0 | 0 | 0 | 908,764 | 0.02 |
| Simple repeat | 0 | 0 | 0 | 0 | 360,628,649 | 6.33 | 360,628,649 | 6.33 |
| Unknown | 0 | 0 | 0 | 0 | 506,285,926 | 8.89 | 506,285,926 | 8.89 |
| Total | 89,667,3423 | 15.75 | 294,809,500 | 5.18 | 2,536,051,111 | 44.53 | 3,069,530,109 | 53.90 |

**Supplementary Table S9** Transposable elements information for the genome of *Sthenoteuthis* sp.

| | Repbase TEs | | TE proteins | | De novo | | Combined TEs | |
|---|---|---|---|---|---|---|---|---|
| Type | Length (Bp) | Percent in genome | Length (Bp) | Percent in genome | Length (Bp) | Percent in genome | Length (Bp) | Percent in genome |
| DNA | 391,738,501 | 6.93 | 4267,081 | 0.08 | 640,298,298 | 11.33 | 798,296,744 | 14.13 |
| LINE | 275,255,583 | 4.87 | 204,179,672 | 3.61 | 424,697,601 | 7.52 | 565,292,718 | 10.00 |
| SINE | 6,468,219 | 0.11 | 0 | 0 | 17,205,635 | 0.30 | 23,645,088 | 0.42 |
| LTR | 99,762,736 | 1.77 | 20,082,106 | 0.36 | 65,053,932 | 1.15 | 153,438,658 | 2.72 |
| Other | 753,396 | 0.01 | 0 | 0 | 0 | 0 | 753,396 | 0.01 |
| Simple repeat | 0 | 0 | 0 | 0 | 14,611,724 | 25.86 | 14,611,724 | 25.86 |
| Unknown | 0 | 0 | 0 | 0 | 1,294,749,419 | 22.91 | 1,294,749,419 | 22.91 |
| Total | 601,351,127 | 10.64 | 228,510,191 | 4.04 | 2,278,563,325 | 40.32 | 2,394,387,829 | 42.37 |

**Supplementary Table S10** 66 positively selected genes for the purpleback flying squids' clades with $P$ <0.01.

| Gene ID in *S.oualaniensis* | Gene ID in *Sthenoteuthis sp.* | Gene name | *P* value |
|---|---|---|---|
| Stoua_s03870_0001 | Ssto026870 | *syncrip* | 0 |
| Stoua_s02768_0003 | Ssto026306 | *atm* | 0 |
| Stoua_s00320_0007 | Ssto024424 | *bmp4* | 0 |
| Stoua_s03733_0006 | Ssto024280 | *xpo7* | 0 |
| Stoua_s01866_0007 | Ssto020258 | *sf3a3* | 0 |
| Stoua_s00815_0001 | Ssto013605 | *nfs1* | 0 |
| Stoua_s00407_0007 | Ssto004645 | *cachd1* | 0 |
| Stoua_s00590_0007 | Ssto014662 | *ars2* | 0 |
| Stoua_s06214_0003 | Ssto011351 | *unc80* | 0 |
| Stoua_s00265_0007 | Ssto027203 | *ascc3* | 0 |
| Stoua_s00393_0008 | Ssto004174 | *prmt7* | 0 |
| Stoua_s02684_0001 | Ssto002215 | *mgea5* | 2.22E-16 |
| Stoua_s06242_0001 | Ssto013541 | *nedd4* | 2.22E-16 |
| Stoua_s00282_0004 | Ssto024669 | *ippk* | 1.55E-15 |
| Stoua_s00564_0008 | Ssto014605 | *ambra1* | 1.55E-15 |
| Stoua_s06540_0001 | Ssto000868 | *ino80d* | 5.33E-15 |
| Stoua_s09115_0002 | Ssto022041 | *znf839* | 5.55E-15 |
| Stoua_s02772_0005 | Ssto027163 | *rabgap1* | 5.02E-14 |
| Stoua_s00009_0041 | Ssto002933 | *dpys* | 1.86E-13 |
| Stoua_s01148_0003 | Ssto025284 | *usp9x* | 8.87E-13 |
| Stoua_s04416_0003 | Ssto008192 | *insig2* | 2.28E-12 |
| Stoua_s01214_0001 | Ssto006769 | *bora* | 2.74E-12 |
| Stoua_s09069_0001 | Ssto007035 | *zfyve26* | 4.80E-12 |
| Stoua_s02175_0003 | Ssto024903 | *mapre1* | 7.10E-10 |
| Stoua_s05524_0002 | Ssto027228 | *fgfr1op* | 7.17E-10 |
| Stoua_s00009_0044 | Ssto005062 | *cd164* | 2.45E-09 |
| Stoua_s10779_0002 | Ssto015283 | *slc25a32* | 5.63E-09 |
| Stoua_s08935_0001 | Ssto025463 | *wdr3* | 1.34E-08 |
| Stoua_s06966_0009 | Ssto004631 | *dtl* | 1.97E-08 |
| Stoua_s02706_0004 | Ssto020642 | *mmaa* | 2.81E-08 |
| Stoua_s00510_0002 | Ssto002166 | *c7orf26* | 6.76E-08 |
| Stoua_s01049_0001 | Ssto021889 | *insr* | 7.11E-08 |
| Stoua_s03610_0005 | Ssto017106 | *ndufa6* | 1.39E-07 |
| Stoua_s01870_0004 | Ssto005171 | *mnt* | 1.62E-07 |
| Stoua_s01367_0006 | Ssto001318 | *ndufs7* | 4.10E-07 |
| Stoua_s20486_0001 | Ssto004173 | *timm17b* | 5.84E-07 |
| Stoua_s03357_0002 | Ssto025730 | *tjp1* | 1.10E-06 |
| Stoua_s00368_0001 | Ssto026187 | *lasp* | 2.73E-06 |
| Stoua_s00185_0019 | Ssto012363 | *pacs2* | 3.57E-06 |
| Stoua_s01496_0003 | Ssto006657 | *fam117b* | 7.23E-06 |
| Stoua_s03049_0002 | Ssto018770 | *mms22l* | 1.03E-05 |

| | | | |
|---|---|---|---|
| Stoua_s00289_0006 | Ssto015278 | *gphn* | 1.92E-05 |
| Stoua_s16128_0001 | Ssto021882 | *chmp1a* | 5.32E-05 |
| Stoua_s06687_0004 | Ssto020141 | *lamtor2* | 5.89E-05 |
| Stoua_s00098_0009 | Ssto025582 | *eef2k* | 6.35E-05 |
| Stoua_s00446_0004 | Ssto015057 | *exosc7* | 6.54E-05 |
| Stoua_s00274_0006 | Ssto015772 | *anapc5* | 6.96E-05 |
| Stoua_s04160_0002 | Ssto002098 | *srrd* | 7.78E-05 |
| Stoua_s02233_0003 | Ssto002765 | *gcc1* | 0.000119295 |
| Stoua_s14705_0001 | Ssto023421 | *znf280c* | 0.000150344 |
| Stoua_s00560_0006 | Ssto020147 | *copa* | 0.000185983 |
| Stoua_s08936_0001 | Ssto008748 | *sf3b3* | 0.000213642 |
| Stoua_s01367_0008 | Ssto001321 | *bzw2* | 0.000500633 |
| Stoua_s01836_0009 | Ssto019519 | *lsm5* | 0.000566332 |
| Stoua_s01685_0005 | Ssto003270 | *none* | 0.001110618 |
| Stoua_s01815_0003 | Ssto000776 | *dnpep* | 0.001140785 |
| Stoua_s02490_0003 | Ssto005680 | *ufl1* | 0.001436811 |
| Stoua_s00848_0004 | Ssto019063 | *tnip2* | 0.001614129 |
| Stoua_s01722_0006 | Ssto019849 | *pgk1* | 0.001957393 |
| Stoua_s00023_0014 | Ssto003187 | *sli* | 0.002458533 |
| Stoua_s10549_0001 | Ssto024230 | *auh* | 0.002882639 |
| Stoua_s21342_0001 | Ssto022797 | *paqr3* | 0.003196648 |
| Stoua_s04858_0002 | Ssto015809 | *pgs1* | 0.003559868 |
| Stoua_s01374_0001 | Ssto001510 | *bsk* | 0.006474727 |
| Stoua_s00354_0012 | Ssto025693 | *calr* | 0.006717453 |
| Stoua_s13358_0001 | Ssto025302 | None | 0.006969673 |

**Supplementary Table S11** Terms from the Function Ontology with FDR < 0.05 for the 66 PSGs of the two *Sthenoteuthis* lineage. The top 20 terms were listed.

| GO ID | GO term | Cluster frequency | Genome frequency of use | FDR |
|---|---|---|---|---|
| phosphoglycerate kinase activity | GO:0004618 | 2 out of 83 genes | 2 out of 28898 genes | < 1.00e-4 |
| phosphomannomutase activity | GO:0004615 | 2 out of 83 genes | 2 out of 28898 genes | < 1.00e-4 |
| elongation factor-2 kinase activity | GO:0004686 | 2 out of 83 genes | 2 out of 28898 genes | < 1.00e-4 |
| cysteine desulfurase activity | GO:0031071 | 2 out of 83 genes | 2 out of 28898 genes | < 1.00e-4 |
| inositol pentakisphosphate 2-kinase activity | GO:0035299 | 2 out of 83 genes | 2 out of 28898 genes | < 1.00e-4 |
| P-P-bond-hydrolysis-driven protein transmembrane transporter activity | GO:0015450 | 2 out of 83 genes | 5 out of 28898 genes | < 1.00e-4 |
| phosphotransferase activity, carboxyl group as acceptor | GO:0016774 | 2 out of 83 genes | 5 out of 28898 genes | < 1.00e-4 |
| protein transmembrane transporter activity | GO:0008320 | 2 out of 83 genes | 8 out of 28898 genes | < 1.00e-4 |
| peptide transmembrane transporter activity | GO:1904680 | 2 out of 83 genes | 8 out of 28898 genes | < 1.00e-4 |
| macromolecule transmembrane transporter activity | GO:0022884 | 2 out of 83 genes | 10 out of 28898 genes | 0.0018 |
| phosphatidylinositol-3-phosphate binding | GO:0032266 | 2 out of 83 genes | 9 out of 28898 genes | 0.002 |
| amide transmembrane transporter activity | GO:0042887 | 2 out of 83 genes | 12 out of 28898 genes | 0.0029 |
| insulin-like growth factor binding | GO:0005520 | 2 out of 83 genes | 11 out of 28898 genes | 0.0031 |
| calmodulin-dependent protein kinase activity | GO:0004683 | 2 out of 83 genes | 11 out of 28898 genes | 0.0033 |
| transferase activity, transferring phosphorus-containing groups | GO:0016772 | 14 out of 83 genes | 1842 out of 28898 genes | 0.004 |
| protein serine/threonine kinase activity | GO:0004674 | 6 out of 83 genes | 390 out of 28898 genes | 0.0094 |
| phosphatidylinositol | GO:1901981 | 2 out of 83 | 15 out of 28898 | 0.010 |

| | | | | |
|---|---|---|---|---|
| phosphate binding | | genes | genes | |
| quinone binding | GO:0048038 | 2 out of 83 genes | 19 out of 28898 genes | 0.011 |
| growth factor binding | GO:0019838 | 2 out of 83 genes | 18 out of 28898 genes | 0.012 |
| kinase activity | GO:0016301 | 12 out of 83 genes | 1524 out of 28898 genes | 0.012 |

**Supplementary Table S12** Terms from the Process Ontology with FDR < 0.05 for the 66 PSGs of the two *Sthenoteuthis* lineage. The top 20 terms were listed.

| GO ID | GO term | Cluster frequency | Genome frequency of use | FDR |
|---|---|---|---|---|
| [2Fe-2S] cluster assembly | GO:0044571 | 2 out of 59 genes | 2 out of 19047 genes | < 1.00e-4 |
| GDP-mannose biosynthetic process | GO:0009298 | 2 out of 59 genes | 2 out of 19047 genes | < 1.00e-4 |
| molybdopterin cofactor biosynthetic process | GO:0032324 | 2 out of 59 genes | 2 out of 19047 genes | < 1.00e-4 |
| replicative senescence | GO:0090399 | 2 out of 59 genes | 2 out of 19047 genes | < 1.00e-4 |
| aging | GO:0007568 | 2 out of 59 genes | 5 out of 19047 genes | 0.013 |
| GDP-mannose metabolic process | GO:0019673 | 2 out of 59 genes | 3 out of 19047 genes | 0.013 |
| cytokinesis | GO:0000910 | 3 out of 59 genes | 26 out of 19047 genes | 0.014 |
| histone phosphorylation | GO:0016572 | 2 out of 59 genes | 6 out of 19047 genes | 0.016 |
| cell aging | GO:0007569 | 2 out of 59 genes | 3 out of 19047 genes | 0.016 |
| intracellular protein transport | GO:0006886 | 7 out of 59 genes | 418 out of 19047 genes | 0.026 |
| response to abiotic stimulus | GO:0009628 | 3 out of 59 genes | 48 out of 19047 genes | 0.031 |
| nucleotide-sugar biosynthetic process | GO:0009226 | 2 out of 59 genes | 11 out of 19047 genes | 0.035 |
| microtubule anchoring | GO:0034453 | 2 out of 59 genes | 13 out of 19047 genes | 0.036 |
| cellular macromolecule localization | GO:0070727 | 7 out of 59 genes | 525 out of 19047 genes | 0.038 |
| protein import into mitochondrial matrix | GO:0030150 | 2 out of 59 genes | 17 out of 19047 genes | 0.038 |
| response to ionizing radiation | GO:0010212 | 2 out of 59 genes | 13 out of 19047 genes | 0.039 |
| cellular protein localization | GO:0034613 | 7 out of 59 genes | 525 out of 19047 genes | 0.040 |
| nucleotide-sugar metabolic process | GO:0009225 | 2 out of 59 genes | 17 out of 19047 genes | 0.040 |
| prosthetic group metabolic process | GO:0051189 | 2 out of 59 genes | 16 out of 19047 genes | 0.042 |

| | | | | |
|---|---|---|---|---|
| molybdopterin cofactor metabolic process | GO:0043545 | 2 out of 59 genes | 16 out of 19047 genes | 0.044 |
